# The Structure of Cilium Inner Junctions Revealed by Electron Cryo-tomography

**DOI:** 10.1101/2024.09.09.612100

**Authors:** Sam Li, Jose-Jesus Fernandez, Marisa D. Ruehle, Rachel A. Howard-Till, Amy Fabritius, Chad G. Pearson, David A. Agard, Mark E. Winey

**Author notes:** Correspondence: Sam Li David A. Agard Mark Winey.

## Abstract

The cilium is a microtubule-based organelle critical for many cellular functions. Its assembly initiates at a basal body and continues as an axoneme that projects out of the cell to form a functional cilium. This assembly process is tightly regulated. However, our knowledge of the molecular architecture and the mechanism of assembly is limited. By applying electron cryo-tomography and subtomogram averaging, we obtained subnanometer resolution structures of the inner junction in three distinct regions of the cilium: the proximal region of the basal body, the central core of the basal body, and the flagellar axoneme. The structures allowed us to identify several basal body and axoneme components. While a few proteins are distributed throughout the entire length of the organelle, many are restricted to particular regions of the cilium, forming intricate local interaction networks and bolstering local structural stability. Finally, by knocking out a critical basal body inner junction component Poc1, we found the triplet MT was destabilized, resulting in a defective structure. Surprisingly, several axoneme-specific components were found to “infiltrate” into the mutant basal body. Our findings provide molecular insight into cilium assembly at its inner junctions, underscoring its precise spatial regulation.

## Introduction

The cilium or flagellum is a hair-like organelle that fulfills many cellular functions, ranging from cell motility to cellular signaling. The cilium assembly initiates when a barrel-shaped basal body (BB) docks underneath the cell membrane and serves as a template for cilium formation. This is followed by building an axoneme at the distal end of the BB, which elongates and projects from the cell surface until it reaches a certain length. Hundreds of proteins are directly involved in the cilium assembly process, which is highly regulated. Mutation of the ciliary components or dysregulation of the assembly process leads to defective cilia with complex phenotypes. These are manifested in humans as ciliopathies, a diverse spectrum of diseases such as microcephaly and primary ciliary dyskinesia (Reiter and Leroux, 2017; Mill et al., 2023).

Cilium assembly is an evolutionarily conserved process (Carvalho-Santos et al., 2010, 2011; Cavalier-Smith, 2014; Jana, 2021). Previous studies show that the centriole and BB exhibit longitudinal ultrastructural variations that can be demarcated into several regions, namely, the proximal, the central core, and the distal regions (Allen, 1969; Geimer and Melkonian, 2004; Greenan et al., 2018; Li et al., 2019; LeGuennec et al., 2021; Zhang et al., 2024; Laporte et al., 2024; Ruehle et al., 2024). In *Tetrahymena*, a new BB first emerges orthogonal to the proximal end of the old BB. This is marked by a tubular hub at the center of the BB with nine spokes radially emanating from the hub. Together, they form a cartwheel scaffold that defines the organelle’s overall 9-fold symmetry (Breugel et al., 2011; Kitagawa et al., 2011; Noga et al., 2022). At the tip of each spoke, a pinhead structure connects the cartwheel to the triplet microtubule (TMT). The TMT comprises a complete A-tubule with 13 protofilaments (pfs) and partial B- and C-tubules that share pfs with the neighboring tubules. Each TMT is linked to its neighbor TMT by a structure called an A-C linker. This proximal region of the BB longitudinally spans ∼150 nm (Figure 1A). In the central core region, the cartwheel terminates and is replaced by an inner scaffold (Figure 1B), a conserved sheath-like structure at the inner circumference of the BB barrel. The inner scaffold is critical for overall BB structural stability and resistance to the external force exerted onto the BB during cilium beating (Guennec et al., 2020; Ruehle et al., 2024). The central core region spans about 300 nm longitudinally and is followed by the distal region that longitudinally spans ∼110 nm. Here, the BB anchors to the plasma membrane and the TMT becomes the microtubule doublet (DMT), where the C-tubules terminate while the A- and B-tubules continue extension. At the distal end of the BB, a specialized compartment called the transition zone marks the transition of the BB to the ciliary axoneme.

**Figure 1.**
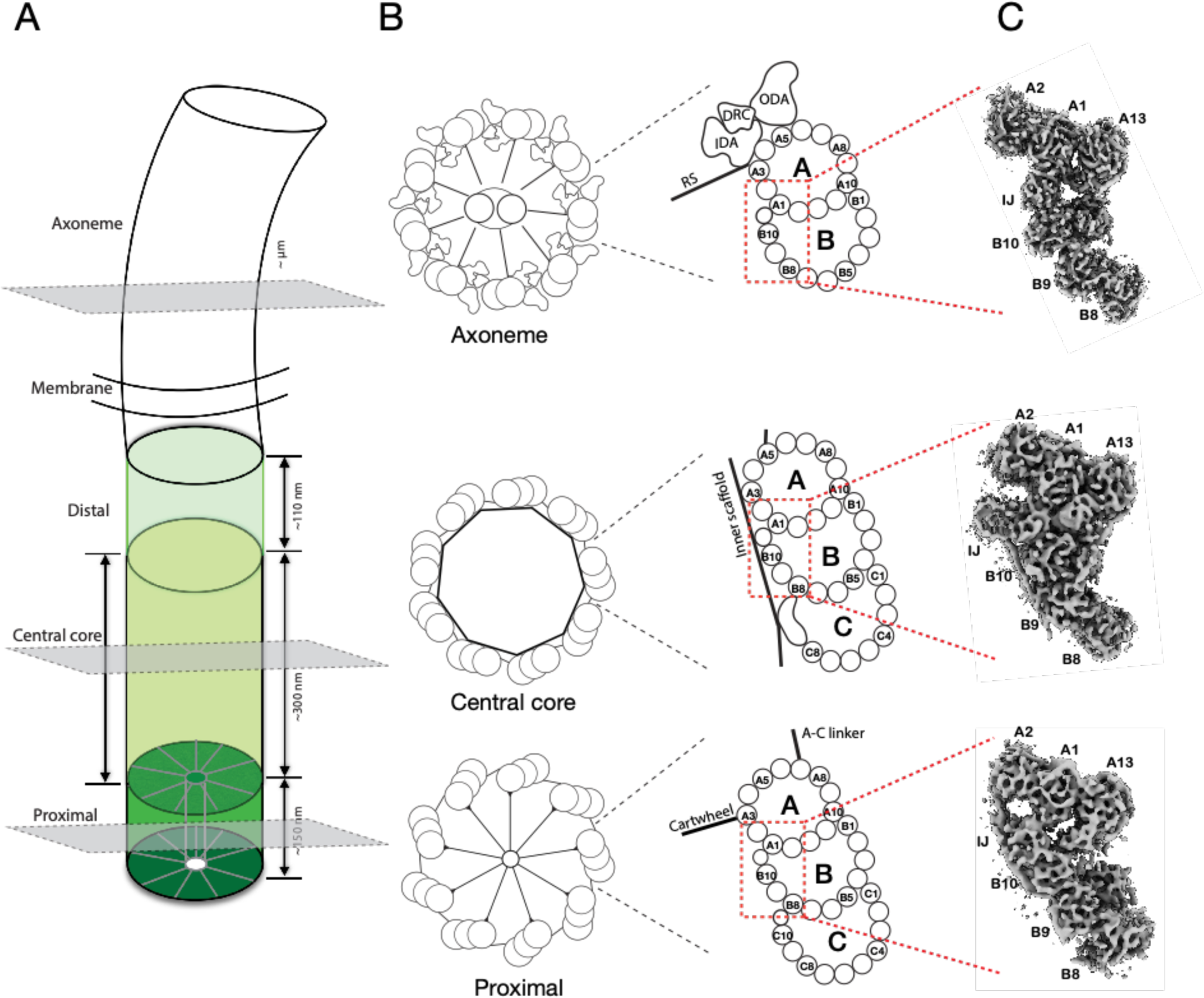
Electron cryo-tomography structures of cilium inner junctions. (A) A schematic diagram of a cilium in *Tetrahymena*, including BB and axoneme. The three regions in the BB, the proximal, central core, and distal, are highlighted in different green colors. Their approximate longitudinal spans are indicated. The three grey-colored cross-sections indicate the location of the structures presented in this work. (B) Left, schematic diagrams of the cross-section of the proximal, the central core region of the BB, and the axoneme. Right, representation of the triplet MT or doublet MT from the three regions. Distinct structures attached to the microtubule wall in each region, such as the cartwheel and A-C linker in the proximal region, the inner scaffold in the central core region, and the Dynein complexes (outer Dynein arm, ODA; inner Dynein arm, IDA; Dynein regulatory complex, DRC) and radial spokes (RS) in the axoneme, are indicated. The red dashed-line boxes highlight the A-B inner junctions (IJ). (C) Three representative subtomograms-averaged structures presented in this work. From bottom to top are the proximal, the central core region of the BB, and the axoneme, as indicated in the cartoons in (B).

The morphological differences in BB ultrastructure indicate variation in its composition along the length of the BB and axoneme. Immuno-EM, super-resolution light microscopy and, in particular, recent advances in Ultrastructure Expansion Microscopy (UExM) have greatly enriched our knowledge of the molecular composition and organization of the BB (Pearson et al., 2009; Hamel et al., 2017; Guennec et al., 2020; Tian et al., 2021; Steib et al., 2020; Arslanhan et al., 2023; Laporte et al., 2024). These studies show that the different BB regions have unique sets of proteins. These composition specificities imply the existence of a spatiotemporal control mechanism that governs the BB assembly. However, the molecular architecture of the BB, the precise boundaries of the composition changes, and the mechanism governing the BB assembly need to be better understood.

The inner and outer junctions, where the B- or C-tubule joins to the neighboring A- or B-tubule, respectively, are unique structures in the TMT and DMT. The inner junction faces the luminal side of the BB, whereas the outer junction faces the exterior of the BB (Figure 1B). Recent studies show that the axonemal inner junction is a protein interaction hub in motile cilia, playing a critical role in withstanding substantial force and stress during cilia beating (Owa et al., 2019; Khalifa et al., 2020). Therefore, it is not surprising that many microtubule inner proteins (MIPs) identified in the axoneme inner junction, such as FAP20, PACRG, FAP52/WDR16, FAP106/ENKUR, FAP45, and FAP210, are evolutionarily conserved across many species (Gui et al., 2021; Kubo et al., 2023). Mutations in many of their human orthologs cause ciliopathies (Zhou et al., 2023; Walton et al., 2022; Gui et al., 2021; Ma et al., 2019; Leung et al., 2023; Sigg et al., 2017; Kubo et al., 2023). However, in contrast to our rich knowledge about the axoneme inner junction structures, little is known about the inner junction in the BB. This limits our understanding of its molecular composition and the mechanistic details of its assembly.

Recently, we identified Poc1 as a key component in the inner junctions critical for BB structure stability and integrity (Ruehle et al., 2024). Here, using cryogenic electron tomography (cryoET) and subtomogram averaging, we extended the study of basal bodies and axonemes isolated from wild-type and mutant strains of a unicellular model organism, *Tetrahymena thermophila*. Based on the inner junction structures in subnanometer resolution, representing three locations of the cilium that cover both BB and axoneme (Figure 1, Figure S1), we identified several components that are universal along the cilium and proteins that are unique to specific regions. This spatial specificity at the inner junction underpins a mechanism that might be applied to the entire cilium assembly during the organelle’s biogenesis.

## Results

### Structure of the inner junction at the BB proximal region

First, we analyzed the TMT inner junction from the BB’s most proximal ∼150 nm region. At 9.82 Å resolution (Figure 2A, Figure S1A, B), the fold of tubulins and many MIPs could be resolved. This facilitated identifying and fitting their corresponding atomic models into the density maps. In addition to the Poc1 that has recently been identified in the A-B and B-C inner junctions (Ruehle et al., 2024), we identified FAP52 and FAP106 in the A-B inner junction in the proximal region. This suggests that all three components, FAP52, FAP106, and Poc1, are recruited to the TMT at the beginning of BB assembly. Like the axoneme, FAP52 and FAP106 have a longitudinal periodicity of 16 nm (Figure 2A). A comparison of the binding of FAP52 to the MT backbone in the proximal region to the central core region reveals marked differences (Figure 2B). Surprisingly, FAP52 in the proximal region (FAP52_prox_) binds predominantly to pf B9 (Figure 2B). This is in contrast to the core region, where FAP52 (FAP52_core_) is at the lateral interface between pf B9 and B10, interacting with both pf, which resembles the structure in axoneme DMT (Ma et al., 2019; Khalifa et al., 2020).

**Figure 2.**
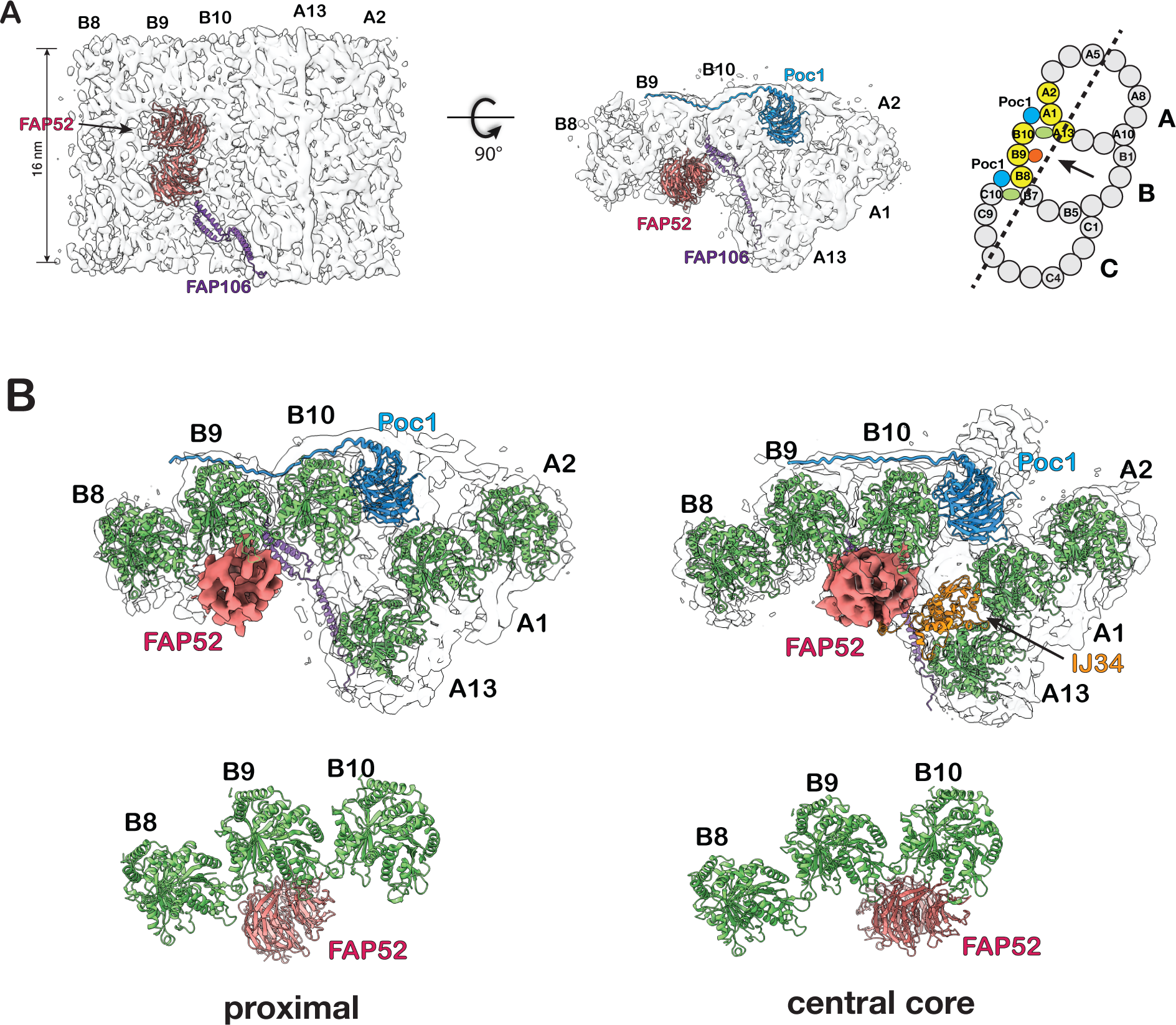
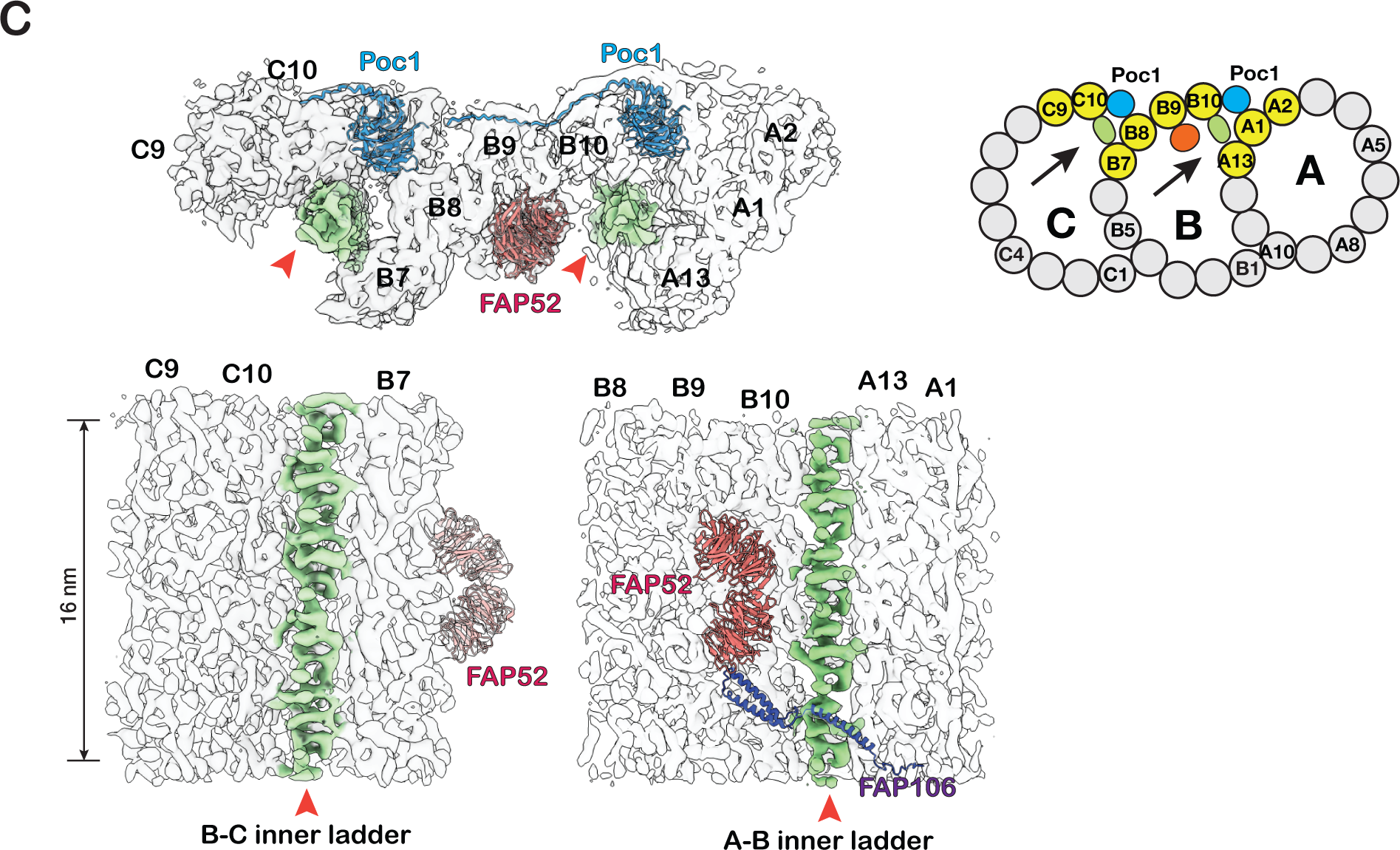
Structure of the inner junctions in the proximal region of the BB. (A) The A-B inner junction and the MIPs are identified in the BB proximal region. A schematic diagram of TMT is on the right. The pfs shown in the structure are highlighted in yellow. A dash line indicates the cutting plane and an arrow indicates the direction of view. (B) Comparison of A-B inner junctions in the BB’s proximal and central core regions. FAP52 binds to B-tubule at different locations in the two structures. The proximal region structures are on the left, and the central core region structures are on the right. (C) Upper: a composite map showing the proximal region’s A-B and B-C inner junctions. The A-B and B-C inner ladders crosslinking pfs A13-B10 and B7-C10 are highlighted green and indicated by red arrowhead. A schematic diagram of TMT is on the right. The pfs shown in the structure are highlighted in yellow. Arrows indicate the directions of view in the bottom panels. Bottom: longitudinal cross-section views of the A-B and B-C inner junctions. The A-B inner ladder and B-C inner ladder are colored green.

In the proximal region, in addition to Poc1, FAP52, and FAP106, we found two previously unidentified structures in the A-B and B-C inner junctions. They connect the A- and B-tubules or B- and C-tubules by crosslinking pf B10 to A13 or C10 to B7, respectively (Figure 2C, Figure S2A). Longitudinally, both structures resemble ladders composed of a stack of rungs. Thus, we named them the A-B inner ladder and B-C inner ladder. Both are unique to the proximal region. Since the A-B inner ladder occupies the luminal side of pf B10, this A-B inner ladder and the FAP52 _core_ at its canonical position between pf B09 and B10 are mutually exclusive.

Interestingly, these A-B and B-C inner ladders remain present in the two inner junction locations in the POC1 knockout mutant (poc1Δ) (Figure S2B), suggesting they have overlapping functions with Poc1 to crosslink the inner junctions and stabilize the TMT in the proximal region. We cannot confirm if these two structures are composed of an identical MIP at our current resolution. It will be interesting to identify these components and study their function in the future.

### The transition from the proximal to the central core region of BB

The finding that FAP52 binds at different locations in the proximal and the core regions implies that the molecule will shift its location near the end of the proximal region before reaching the core region. To investigate this shift in more detail and to identify any additional changes associated with this shift, we applied 3D classification on the subtomograms dataset from the BB proximal region, focusing on the A-B inner junction. The result shows two groups where FAP52 is at different locations (Figure 3A, Figure S3A). A predominant group (Class 1, 6030 subtomograms, 83.2%) shows the FAP52 binding at pf B9, representing the proximal region structure. In a minor group (Class 2, 1218 subtomograms, 16.8%), the FAP52 binds to pf B9/B10, a “canonical” binding location by FAP52 both in the central core region of BB and in the axoneme (Figure 3A, Figure S3A). We backtracked their location in the BB and generated a histogram representing the longitudinal distribution of these two groups (Figure 3B). The distribution histogram shows that the transition occurs approximately at a longitudinal position of 130 nm. This minor group likely represents the structure near the distal end of the proximal region, where FAP52 shifts from pf B9 to its canonical position at pf B9/B10. To visualize this shift, we calculated the average of this minor group in a larger volume, longitudinally spanning 80 nm, including additional structure features towards the proximal end, where it shows the FAP52 binding to pf B9 (Figure 3C, Figure S3B, Movie S1). Thus, this extended volume confirms the change of FAP52 taking place at ∼130 nm.

**Figure 3.**
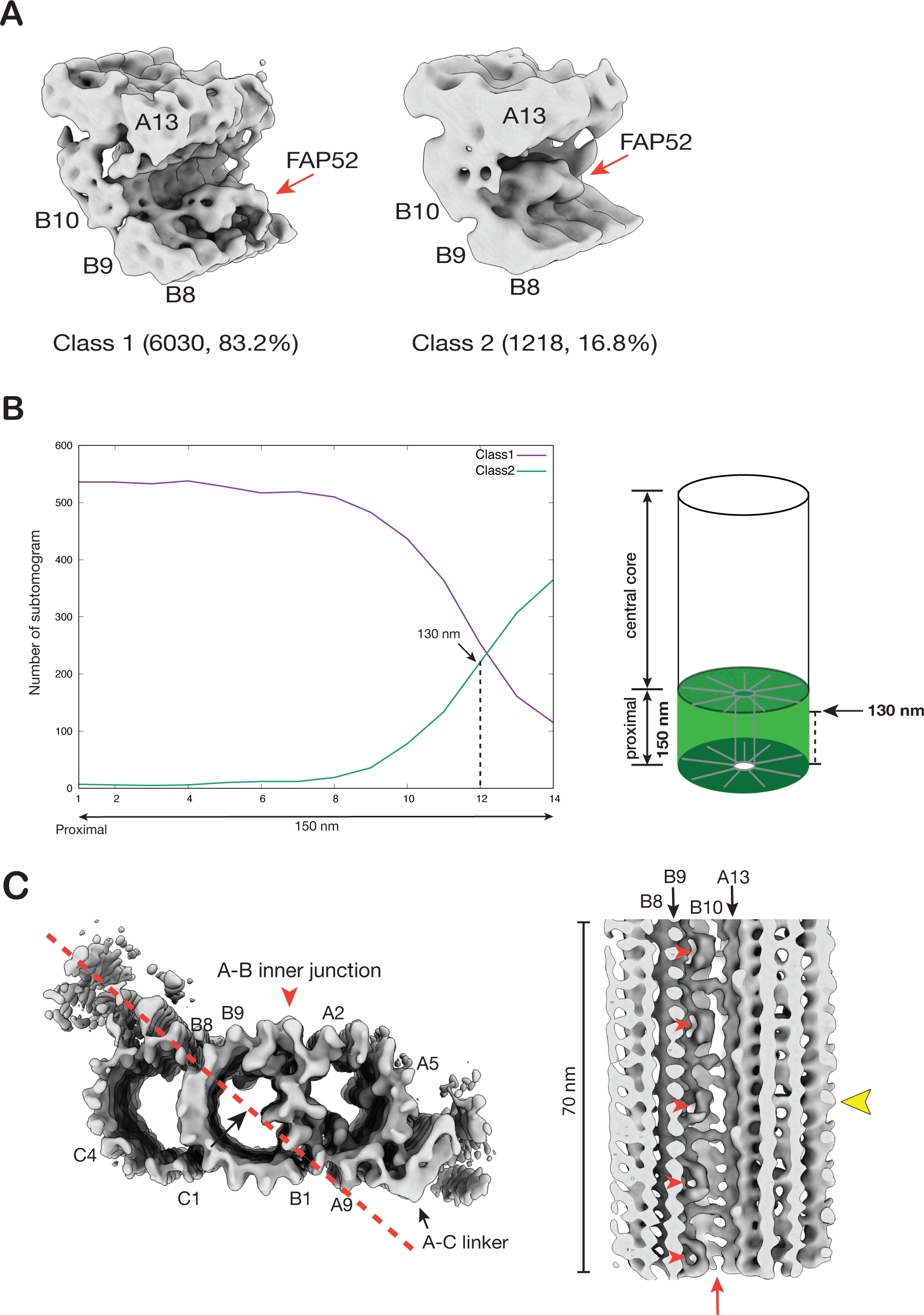
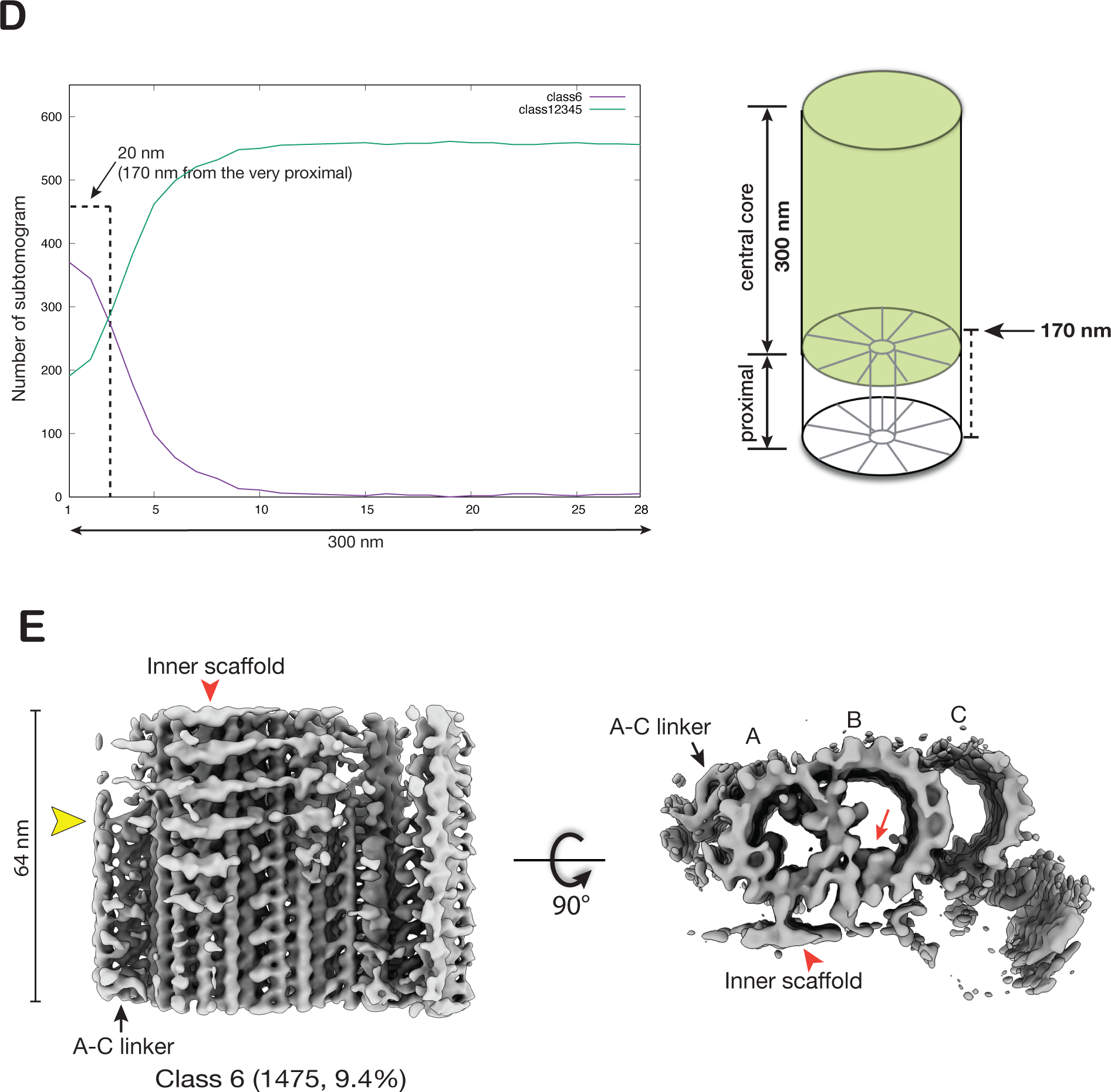
Structural changes at the transition from the proximal to the central core region of the BB. (A) Focused classification on the subtomograms from the proximal region identifies two structures where FAP52 binds at different locations (indicated by red arrows). (B) The longitudinal distribution of the subtomograms from the proximal region. The 150 nm long proximal region is divided into 14 bins. Based on the two classes identified in (A), for each class, the number of subtomograms (y-axis) found in each bin is plotted along the length (x-axis), showing their longitudinal distribution. *Right*: A schematic diagram indicates the weighted average longitudinal length for Class 2 is 130 nm. The weighted average length is L = ∑ j*Nj / ∑ Nj (Nj: number of subtomograms found in bin j). (C) Left: The Class 2 average in a cross-section view. A red arrowhead indicates the A-B inner junction. A red dashed line and a black arrow indicate the cross-section and viewing direction of the structure on the right. *Right*: The Class 2 average longitudinally expanded to 70 nm shows FAP52 (red arrowheads) shifting binding site, indicated by a yellow arrowhead. The shift coincides with the termination of the unidentified ladder-like protein in the A-B inner junction (red arrow). (D) Similar to (B), the longitudinal distribution of the subtomograms from the central core region. The 300 nm long central core region is divided into 28 bins. Based on the classification result shown in (Figure S3C), the longitudinal distribution of subtomograms in Class 6 and the other five classes are plotted. The weighted average longitudinal position of Class 6 is at 170 nm from the very proximal end of the BB. This is illustrated in a schematic diagram on the right. (E) The averaged structure in Class 6 (Figure S3C) shows the changes of TMT transitioning from the proximal to the central core region. A red arrowhead indicates the inner scaffold. A red arrow indicates FAP52. A yellow arrowhead indicates the A-C linker’s termination and the inner scaffold’s emergence.

Interestingly, the shift of FAP52 from its pf B9 position to the canonical pf B9/B10 position is concomitant with the termination of the A-B inner ladder at the inner junction (Figure 3C, Movie S1). Likely, the termination of this A-B inner ladder at 130 nm longitudinal position makes space for FAP52. Furthermore, the A-C linker, a characteristic feature of the proximal region, remains in the structure at this longitudinal position. In contrast, the inner scaffold, a structure unique to the BB central core region, is absent (Figure 3C). This indicates that, at this point, the central core region has yet to be established when FAP52 shifts the location.

To locate and visualize the transition from the proximal to the central core region, we applied 3D classification on the subtomograms dataset from the BB central core region. The classification identified a minor subset (1475 subtomograms, 9.4%) showing both the A-C linker and the inner scaffold (Figure S3C). A longitudinal distribution plot shows that, in contrast to other subsets with subtomograms evenly distributed throughout the central core region, this subset is concentrated at the beginning of the central core region, centered at 170 nm longitudinal position (Figure 3D). A 3D average of this subset shows the A-C linker’s wane and the inner scaffold’s concomitant emergence (Figure 3E, Movie S1). These likely mark the transition from the proximal to the central core region of the BB.

In summary, our structural analysis of the BB’s most proximal 150 nm stretch shows a series of sequential events in the inner junctions. First, we found FAP52, FAP106, and Poc1 in the proximal region of the BB. Surprisingly, FAP52, one of the most conserved inner junction components of BBs and axonemes, binds at pf B9, while two unidentified components forming ladder-like structures crosslink A-B and B-C inner junctions, respectively. Second, at about 130 nm from the proximal end, the inner ladders terminate. This coincides with the FAP52 shifting from pf B9 to a “canonical” position at pf B9/B10. Third, at ∼170 nm, the A-C linker terminates and the inner scaffold emerges, marking the transition from the proximal to the central core region. All these changes occur in well-defined longitudinal positions and are displayed sequentially, demonstrating a tight spatial regulation for the BB MIPs.

### Structure of the inner junction in the core region and comparison to the flagellar axoneme

In *Tetrahymena*, the central core region of BB spans longitudinally about 300 nm, from 150 nm to 450 nm (Figure 1). The A-B inner junction in this region has important structural roles. First, it crosslinks the A- and B-tubule in the luminal side of the BB, stabilizing the TMT. Second, it connects the TMT to the inner scaffold, a unique structure in the core region critical for the overall BB stability and structural cohesion (Guennec et al., 2020; Ruehle et al., 2024). Poc1, a WD40 family protein, occupies the A-B inner junction, providing an anchor that connects the inner scaffold to the TMT. In the axonemal DMT, alternating FAP20 and PACRG filaments fill the A-B inner junction (Ma et al., 2019; Gui et al., 2021; Khalifa et al., 2020; Dymek et al., 2019). Based on the previously solved axoneme structures (Ma et al., 2019; Kubo et al., 2023), we built atomic models into the subnanometer density maps from the BB core and axoneme regions (Figure 4A, 4B). This allowed us to identify several axoneme MIPs in the A-B inner junction of the BB core region. These include FAP52, FAP106, IJ34, FAP45, and FAP210. These MIPs are in the same locations as the axoneme DMT, with the same longitudinal periodicity.

**Figure 4.**
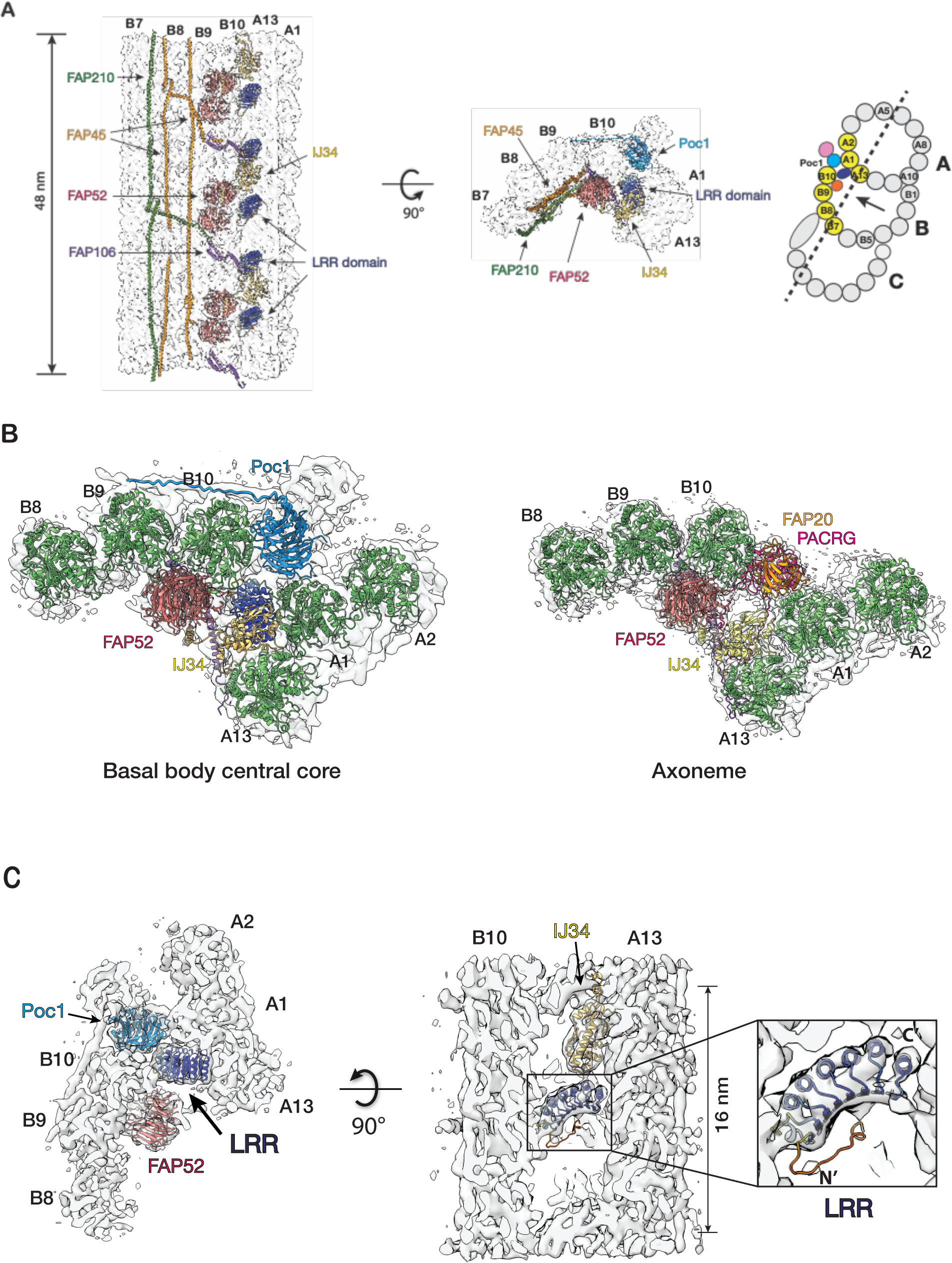
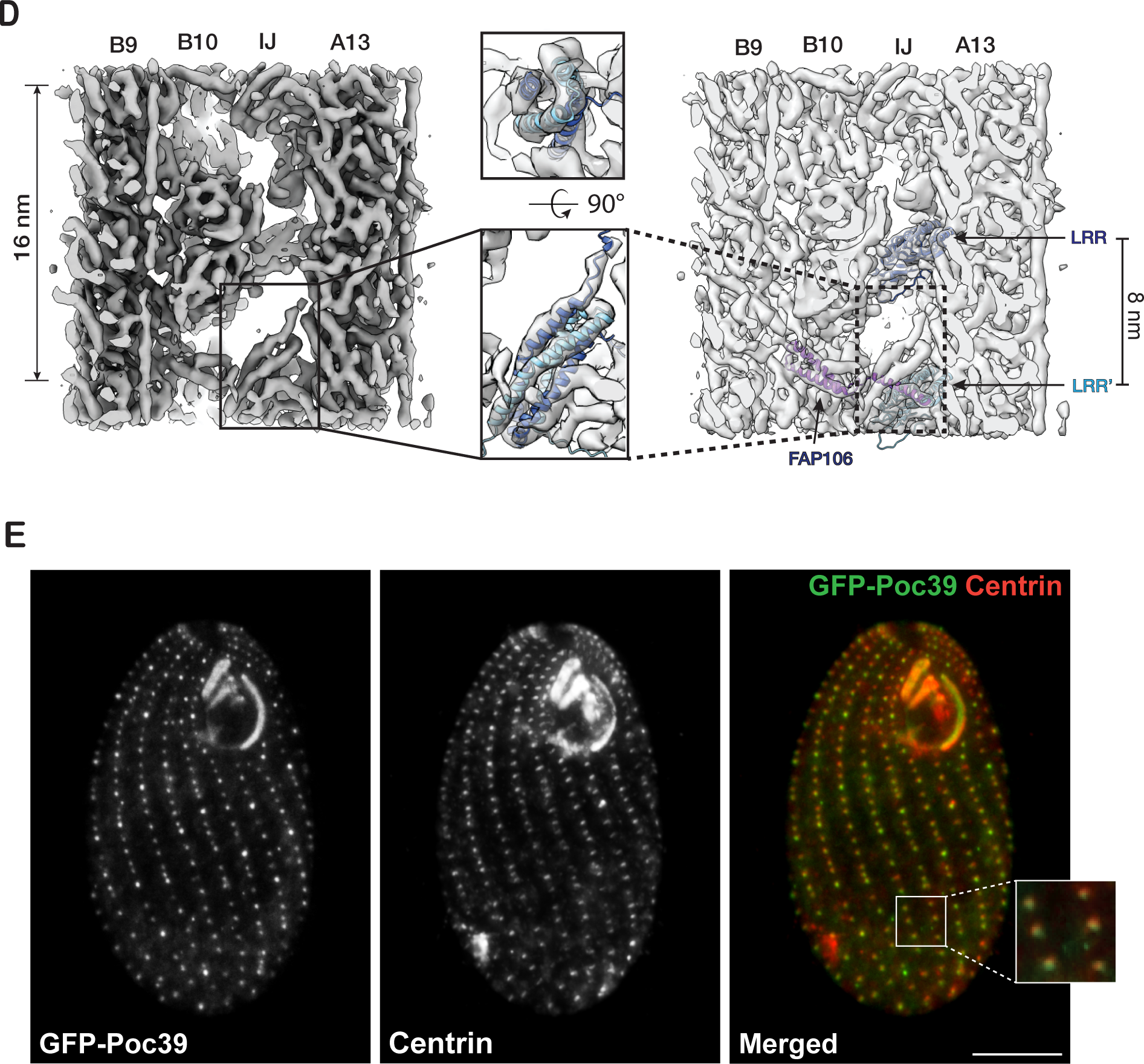
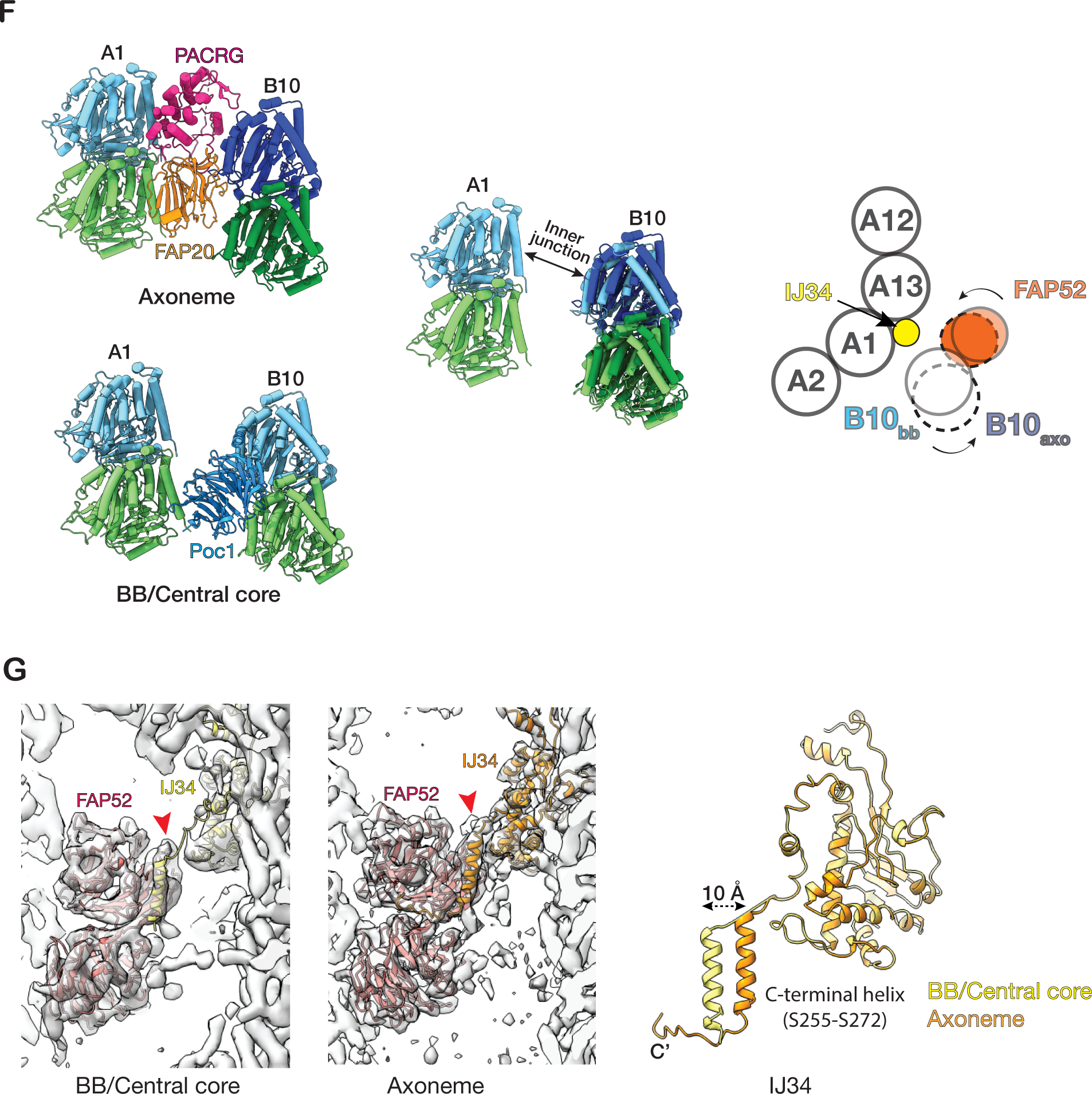
Structure of the inner junctions in the central core region of the BB. (A) Two orthogonal views of the 48-nm repeat structure at the A-B inner junction. The pfs shown in the structure are highlighted in yellow in a schematic diagram of the TMT on the right. A dash line indicates the cutting plane and an arrow indicates the direction of the view. (B) Comparing the A-B inner junction from the BB central core region to the axoneme. (C) A MIP with a leucine-rich repeat (LRR) motif in the A-B inner junction. An LRR model predicted by AlphaFold2 from the protein (UniProt: Q22N53) fits into the density map. The LRR motif makes potential interactions with pfs B10, A13, and A1 of the TMT wall, Poc1, and IJ34. (D) An AlphaFold3 predicted model of a right-handed 4-helix bundle fits into the density map. It is formed by two NTDs from the neighboring LRR-containing proteins as an antiparallel dimer. Each monomer is colored in either dark or light blue. (E) Localization of Poc39 to the *Tetrahymena* BB by fluorescence microscopy. Left: fluorescence signal of GFP-Poc39 expressed in *Tetrahymena* cell. Center: immunofluorescence signal of a BB protein Centrin in the same cell. Right: merge of the two images on the left. The inset is an enlarged view of a local area showing the colocalization of the GFP-Poc39 (green) and the Centrin (red) signals. Scale bar: 10 μm. (F) Comparing the inner junctions between the central core region and the axoneme. Left: the models of the inner junction in the central core of BB and the axoneme. Center: superposition of the two models using pf A1 as a reference. Pf B10 (α/β tubulin) in the BB central core are in light green and blue. Pf B10 (α/β tubulin) in the axoneme are dark green and blue. Right: schematic illustration of the change. The solid circles represent the pf B10 and FAP52 from the central core. The dashed circles represent the pf B10 and FAP52 in the axoneme. The two curved arrows indicate the movement of pf B10 and FAP52 from the central core to the axoneme. (G) Comparing the structures of IJ34 in the central core region and the axoneme. The left and center show that IJ34 and FAP52 fit into the density maps. Right: superposition of the two IJ34 using their main domains as a reference. Their C-terminal helices are 10 Å apart.

In addition to these MIPs in both BBs and axonemes, we identified a MIP unique to the BB core region (Figure 4C, 4D). It crosslinks the A- and B-tubule by binding to pfs A13 and B10 with an 8 nm periodicity. At 8.31 Å resolution (Figure S1D), the map shows an overall double-layer arch-shaped topology (Figure S4A). The top layer comprises at least seven α-helices, whereas the bottom layer is continuous, likely a β-sheet. This tertiary arrangement suggests a leucine-rich repeat (LRR) motif. Searching the BB proteome database identified seven proteins containing LRR motifs (Kilburn et al., 2007). Based on the AI-based structure prediction (Jumper et al., 2021), one of the proteins’ (UniProt Q22N53) C-terminal LRR motif fits well into the density map (Figure 4C, S4B, Movie S3). In the model, the LRR is an arch connecting the A- and B-tubule at the inner junction. Specifically, the N-terminal end of the LRR motif potentially contacts an α-tubulin from pf B10, while at the opposite side of the arch, the C-terminus of the LRR motif binds at a four-tubulin corner formed by pf A13 and A1. In addition, the LRR also makes potential contact with Poc1 and IJ34 (Figure 4C, Movie S3). The N-terminal portion of the protein is predicted to form an anti-parallel hairpin of two α-helices, albeit with low confidence scores (pLDDT) (Figure S4B). Interestingly, our averaged density map observed a right-handed 4-helix bundle between two neighboring LRR-containing proteins (Figure 4D). This 4-helix bundle has 16-nm longitudinal periodicity and makes contact with FAP106. An AlphaFold 3 prediction (Abramson et al., 2024) shows that two copies of the α-helical hairpin could dimerize to form a 4-helix bundle (Figure S4C, S4D). This is further supported by fitting this predicted 4-helix bundle into the density map (Figure 4D), showing an overall good agreement between the prediction and the experimental observation. Next, to localize this LRR-motif-containing protein in the cell, we labeled the protein with GFP and expressed the fusion protein in *Tetrahymena*. The GFP signal colocalized with the BB component Centrin, confirming the protein’s localization at the BB (Figure 4E). Thus, based on its molecular weight, we named this protein Poc39 (Protein of centriole 39). Given the location of Poc39 in the BB and its interactions with neighbors, it likely plays a critical role in stabilizing the inner junction in the central core region. It will be essential to study its function in the future.

While Poc1 and Poc39 occupy the inner junction in the BB central core region, the equivalent position in the axoneme is occupied by PACRG and FAP20 (Figure 4B). This composition difference might lead to local structural alteration. A comparison of the microtubule arrangement in these two regions shows noticeable variation. Using the pf A1 as a reference, the relative position and the orientation of pf B10 change from the BB inner core region to the axoneme (Figure 4F, Movie S4). Compared to the BB core region, in the axoneme, pf B10 has shifted further away from the A-tubule (∼ 6 Å), increasing the gap between pf A1 and pf B10 and enabling it to accommodate a PACRG and FAP20 protofilament. Meanwhile, pf B10 rotates counter-clockwise about 6 degrees (cartoon in Figure 4F, viewed from the MT plus end). This shift and rotation of pf B10 brings its associated FAP52 closer to the A-tubule (cartoon in Figure 4F). Interestingly, these structural changes in the MT lattice and FAP52 can be accommodated by IJ34, one of the inner junction MIPs present in both the BB and axoneme in *Tetrahymena* (Figure 4G). The IJ34’s C-terminal helix (S255-S272) can transverse about 10 Å and remain tethered to FAP52 in both structures, while its main globular domain anchors to the A-tubule (Figure 4G). Likely, this is enabled by a flexible linker in IJ34 connecting the main globular domain to the C-terminal helix tail.

In addition to the structure change on MIPs, we also observed changes in the MT lattice. The lateral inter-protofilament curvature at pf B9/B10 is markedly higher in the core region (31.6°) compared to the proximal (20.1°) or the axoneme region (23.3°) (Figure S4E, Table 2), demonstrating the elasticity of the MT lattice that is capable of forming irregular and highly variable lateral curvatures. This is consistent with previous studies on different forms of chemically treated axoneme structures (Ichikawa et al., 2019), as well as a study on *in vitro* assembled MTs revealing its highly flexible and polymorphic lateral inter-pf interactions (Debs et al., 2020).

**Table 1.**
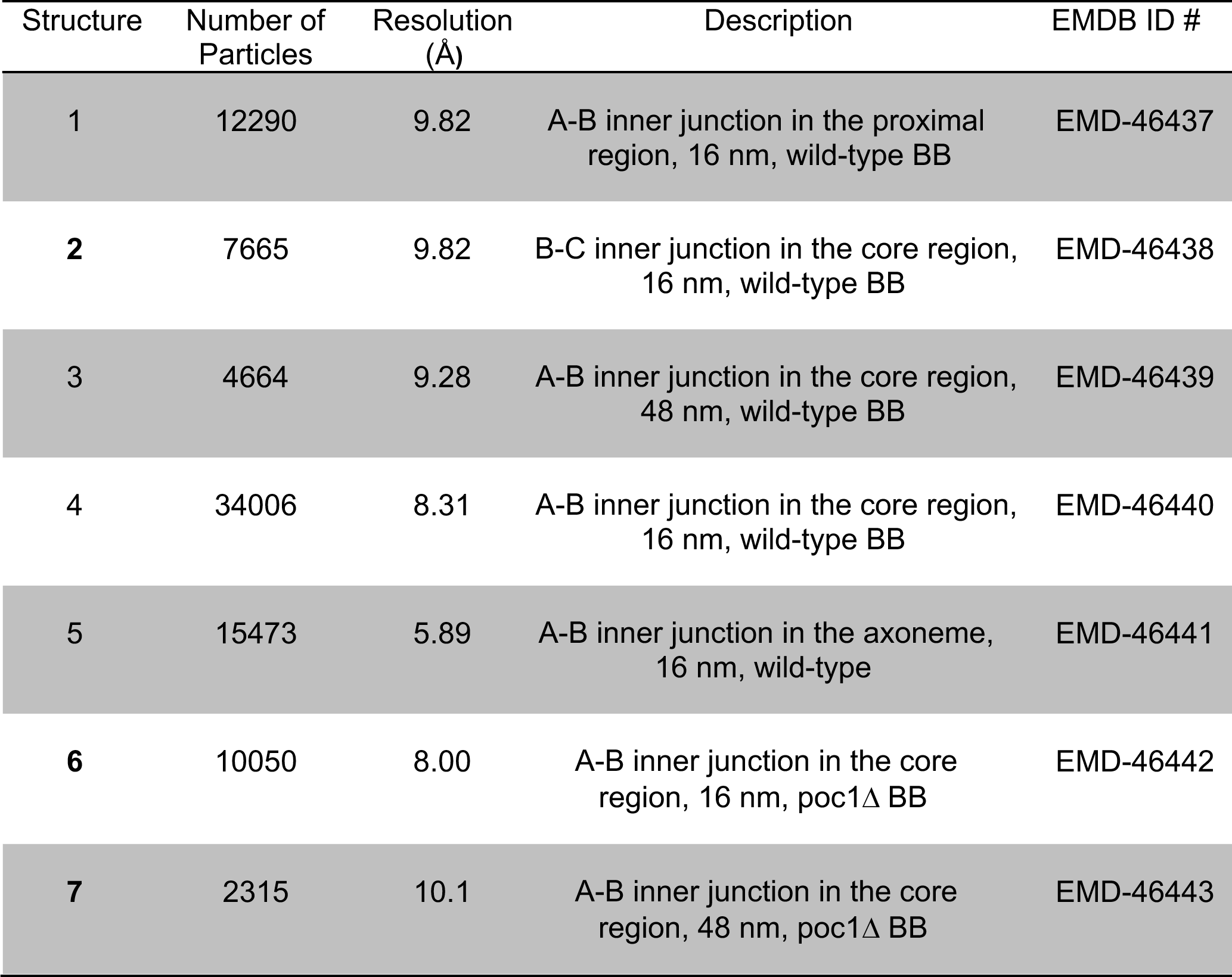
Summary of Structures.

| Structure | Number of Particles | Resolution (Å) | Description | EMDB ID # |
| --- | --- | --- | --- | --- |
| 1 | 12290 | 9.82 | A-B inner junction in the proximal region, 16 nm, wild-type BB | EMD-46437 |
| 2 | 7665 | 9.82 | B-C inner junction in the core region, 16 nm, wild-type BB | EMD-46438 |
| 3 | 4664 | 9.28 | A-B inner junction in the core region, 48 nm, wild-type BB | EMD-46439 |
| 4 | 34006 | 8.31 | A-B inner junction in the core region, 16 nm, wild-type BB | EMD-46440 |
| 5 | 15473 | 5.89 | A-B inner junction in the axoneme, 16 nm, wild-type | EMD-46441 |
| 6 | 10050 | 8.00 | A-B inner junction in the core region, 16 nm, poc1Δ BB | EMD-46442 |
| 7 | 2315 | 10.1 | A-B inner junction in the core region, 48 nm, poc1Δ BB | EMD-46443 |

**Table 2.**
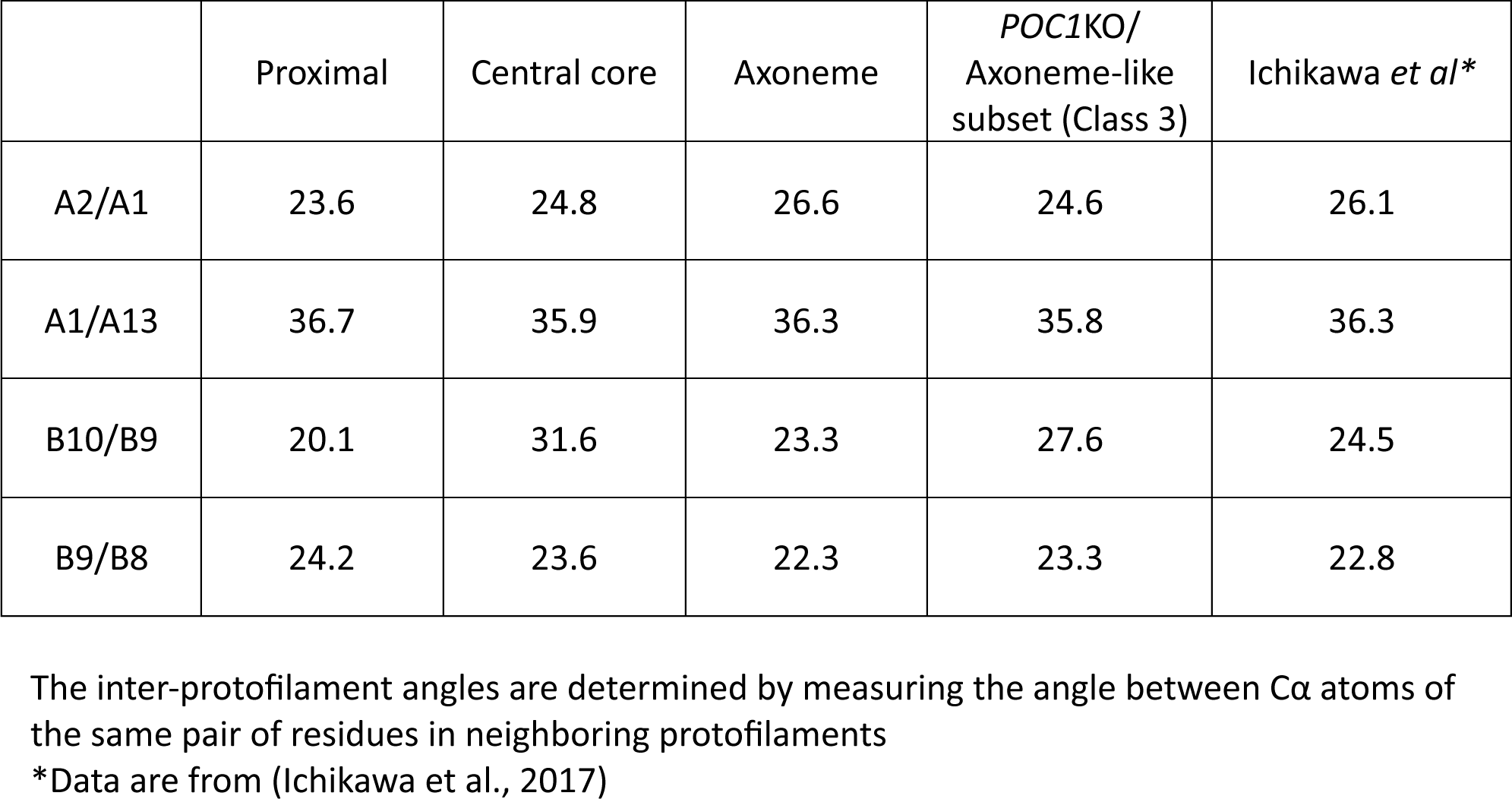
Measurement of inter-protofilament angle (Δφ, in degree)

### Incorporation of a set of axonemal proteins in the BB inner junction in the *POC1* knockout mutants

Recently, we identified Poc1 as a critical component at the inner junctions of TMT (Ruehle et al., 2024). In the proximal region, Poc1 is localized at both the A-B and B-C inner junctions. In the central core region, Poc1 is in the A-B inner junction, providing one of the anchoring sites for the inner scaffold. The poc1Δ shows that the protein plays essential roles in BB stability and resisting external force (Junker et al., 2022; Ruehle et al., 2024). Here, we further characterize the poc1Δ defects in the BBs. Consistent with the previous observations, in poc1Δ, the BB is partially disintegrated, where the B-tubule is partially detached from the A-tubule. However, occasionally, we found that the B-tubule remained attached to the A-tubule at various longitudinal locations (Figure 5A). To investigate this further, we classified the subtomograms from poc1Δ TMT, focusing on the A-B inner junction in the central core region. The classification identified three subsets showing distinct features in the inner junction (Figure 5B). In the first subset (Class 1, 32.5% of the total population), the B-tubule is detached from the A-tubule, resulting in a gap at the inner junction between the A- and B-tubules. This is consistent with our recent observation that Poc1 is critical for BB stability (Ruehle et al., 2024). In the second subset (Class 2, 20.3%), the inner junction remains sealed, albeit without Poc1(Figure 5B). Further refinement of this subset shows that, despite the absence of Poc1 in the inner junction, many MIPs remain, including IJ34, Poc39, FAP106, and FAP52 (Figure 5B, 5F). It is noticeable that in this second subset, while Poc39 remains bound to pf A1/A13 in the A-tubule, it is partially detached from the B-tubule (Figure 5F), indicating a weakened interaction with its neighbors, resulting in a flexible and less well-defined inner junction without Poc1. Unexpectedly, we identified extra proteins in the third subset (Class 3, 47.2%) in the position of Poc1 (Figure 5B). Further refinement of this subset improved the structure to 8.00 Å, resolving the secondary structure of the protein folding (Figure S1F). This allowed us to identify these proteins as PACRG and FAP20 at the A-B inner junction (Figure 5C, Movie S5). Both PACRG and FAP20 are conserved axonemal inner junction components. However, they have not been found in the A-B inner junction in the central core region of the wild-type BB (Ruehle et al., 2024). In addition to PACRG and FAP20, other MIPs IJ34, FAP106, FAP52, FAP45, and FAP210 could also be identified in this subset (Class 3) of the poc1Δ BB (Figure 5E, 5F, S5C). In contrast, Poc39 was absent in this group (Figure 5F). Further classification of this subset identified an additional MIP in 48-nm periodicity previously found unique to the inner junction in wild-type *Tetrahymena* axonemal DMT (Figure 5E, S1G, S5C) (Li et al., 2022). This MIP was proposed to be CCDC81b (Uniprot ID, I7M688) and BMIP1 (Uniprot ID, I7MB72) (Gao et al., 2024). Overlaying this mutant structure (Class 3) with the wild-type axoneme structure shows that these structures are nearly identical (Figure S5A, S5B, RMSD < 1 Å). In contrast, comparing this axonemal-like structure from poc1Δ BB to the wild-type central core region shows noticeable differences (Figure 5D). In the poc1Δ mutant, the pf B10 is shifted (7.1 Å) and rotated (5.5°) relative to the A-tubule, resulting in an increased inner junction gap where the PACRG and FAP20 now fill in. Meanwhile, FAP52 moved closer to the A-tubule, similar to the movement from the central core region to the wild-type axoneme, as described in Figure 4F. In summary, by incorporating axonemal components into the A-B inner junction, a subset (Class 3) of poc1Δ mutant BBs adopted a structure resembling the wild-type axonemal inner junction.

**Figure 5.**
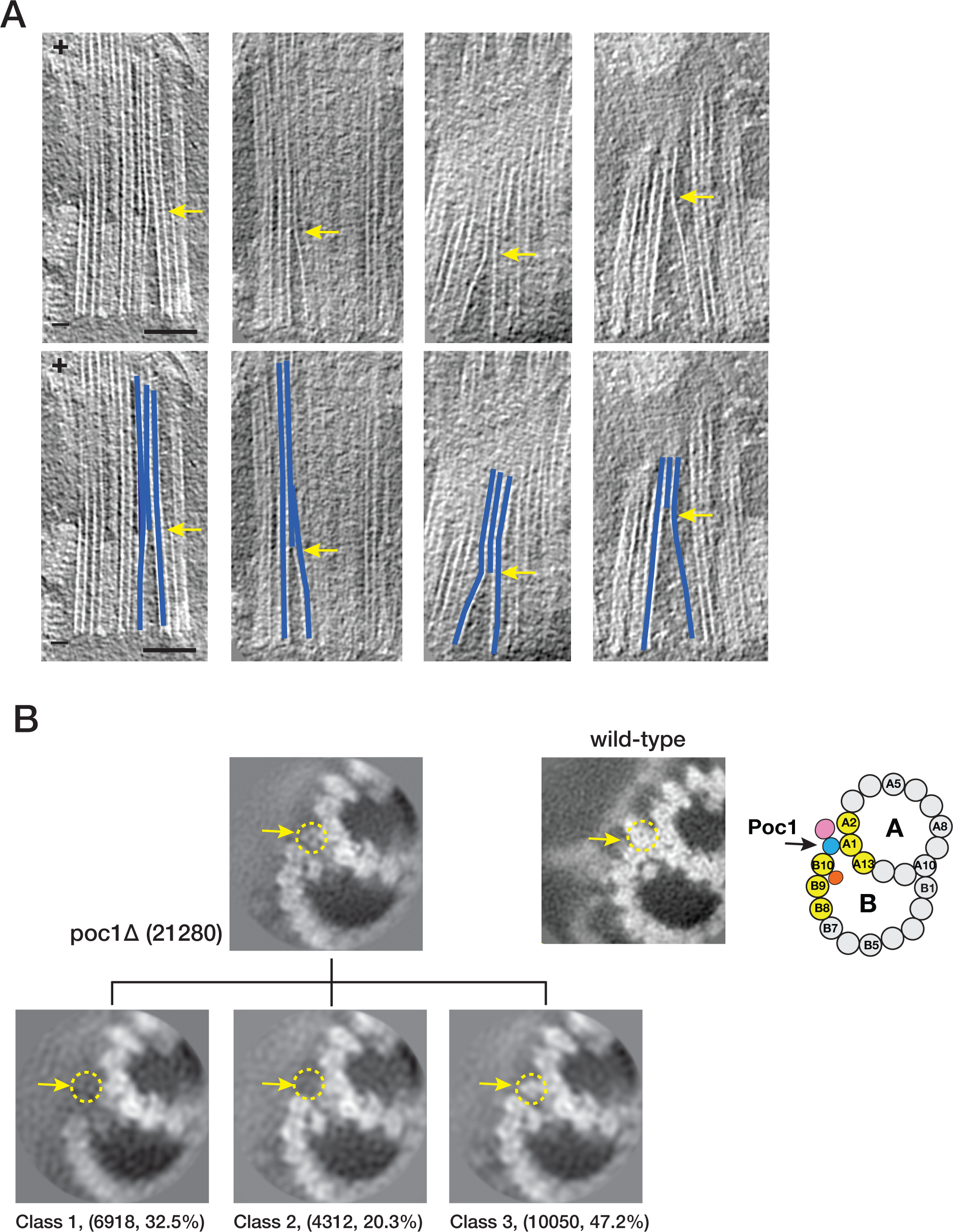
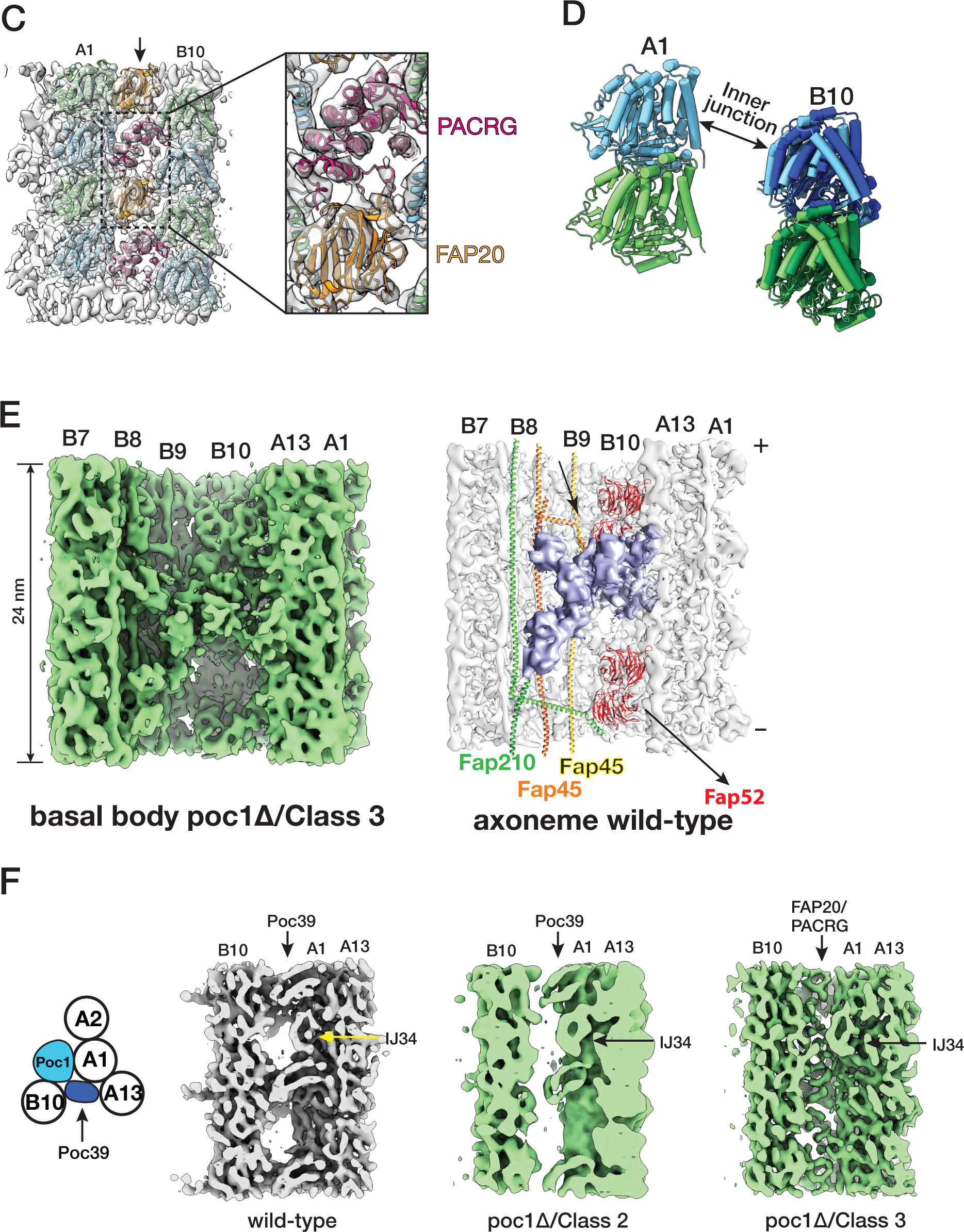
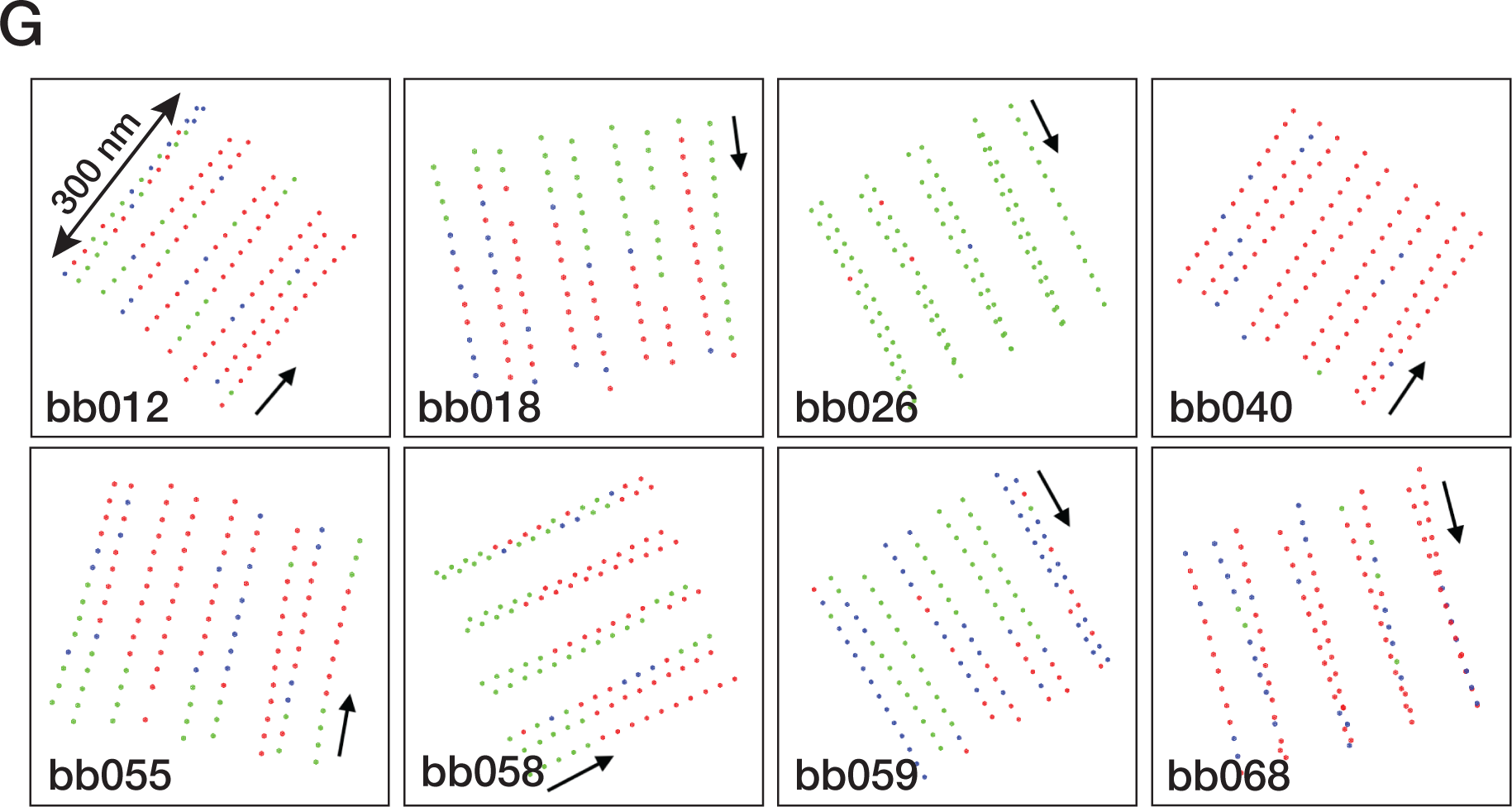
poc1Δ mutants destabilize the A-B inner junction and allow axonemal components to incorporate into the assembly. (A) Upper: representative tomogram slices from poc1Δ BB show the TMTs split at the A-B inner junction at various longitudinal locations. Bottom: the blue lines highlight the split TMTs in the upper panels. The yellow arrows indicate the splits. The minus ends of TMT are at the bottom, and the plus ends are at the top. Scale bar: 100 nm. (B) 3D classification of subtomograms from poc1Δ BB, focusing on the A-B inner junction. The resulting three subsets show structure variations at the A-B inner junction. In Class 1, the B-tubule is incomplete and detached from the A-tubule at the inner junction. In Class 2, the B-tubule is complete and remains attached to the A-tubule, though the Poc1 position is empty. In Class 3, the B-tubule is complete and connected to the A-tubule, while other proteins occupy the Poc1 site. Yellow dashed circles and arrows outline the position of Poc1 in the wild-type. An image from the wild-type and a schematic diagram is shown for comparison. (C) Further refinement of Class 3 at 8.00 Å shows the FAP20/PACRG filament at the mutant’s inner junction gap. (D) Comparing the A-B inner junction in the central core region between wild-type and poc1Δ Class 3 shows an increased gap in the junction. Pf A1 is used as a reference for superposition. For Pf B10 (α/β tubulin), the wild-type is in light green and blue, and the poc1Δ is in dark green and blue. (E) Focused classification of subtomograms from poc1Δ BB (Class 3) identifies additional axoneme MIP. Left, the mutant structure is shown in green. Right, the inner junction from the wild-type axoneme DMT. The corresponding MIP is colored in lavender. (F) In poc1Δ BB, Poc39 partially remains in the Class 2 average but is absent in Class 3, where the inner junction is occupied by FAP20/PACRG. For comparison, the wild-type structure is shown in grey. (G) Mapping the location of subtomogram from 3 subsets identified in the central core region of poc1Δ BBs. 8 representative BBs are shown. The green dots represent the Class 1 subset, an incomplete B-tubule detached from the A-tubule at the inner junction. The blue dots represent the Class 2 subset, complete B-tubule without Poc1. The red dots represent the Class 3 subset, the axoneme-like inner junction where FAP20/PACRG fills in the space left by Poc1. The arrows point in the direction from the proximal to the distal end of the BBs.

Finally, to find if there is any arrangement pattern for these three identified subsets, we mapped them to their corresponding locations in the core region for all 85 poc1Δ BB analyzed (Figure 5G). A consistent pattern was not identified for any of these three subsets. Instead, the distribution shows sporadic dispersion throughout the longitudinal length of the central core region. For example, in one of the BBs (bb026), nearly all subtomograms have incomplete B-tubule at the inner junction. In contrast, in another BB (bb040), almost all inner junctions are found to be “axonemal-like” and were filled by PACRG/FAP20. However, in many BBs, the “axonemal-like” subtomograms aligned continuously, forming a single file in the same TMT, indicating a degree of cooperativity in PACRG/FAP20 assembly into the TMTs. It is conceivable that binding the first PACRG and FAP20 pair is the rate-limiting step, where the local geometry is not optimal for incorporation. However, once they are “wedged” into the inner junction, the local geometry, such as the gap between the pf B10 and A1 and the inter-protofilament curvature, becomes optimal, thereby facilitating recruiting additional PACRG/FAP20 pairs, manifested as cooperativity.

In summary, consistent with our recent study (Ruehle et al., 2024), we found that in the poc1Δ mutant, the assembly at the inner junction is disrupted, showing structural heterogeneity and composition variations. Without Poc1, the inner junction is structurally destabilized and accessible. This allows a set of axoneme components, including PACRG and FAP20, to be incorporated into the BB inner junction, leading to a change of the local MT lattice. These changes likely will propagate and have a long-range impact on the structural integrity of the TMT and BB.

## Discussion

In this work, we use cryoET and image analysis to study the structures of basal bodies and axonemes isolated from *Tetrahymena thermophila*. By focusing on the inner junction at three distinct locations, the proximal region, the central core region of the BB, and the axoneme region, we identified several MIPs present throughout BBs and axonemes and proteins specific to a distinct region. These are summarized in Figure 6A.

**Figure 6.**
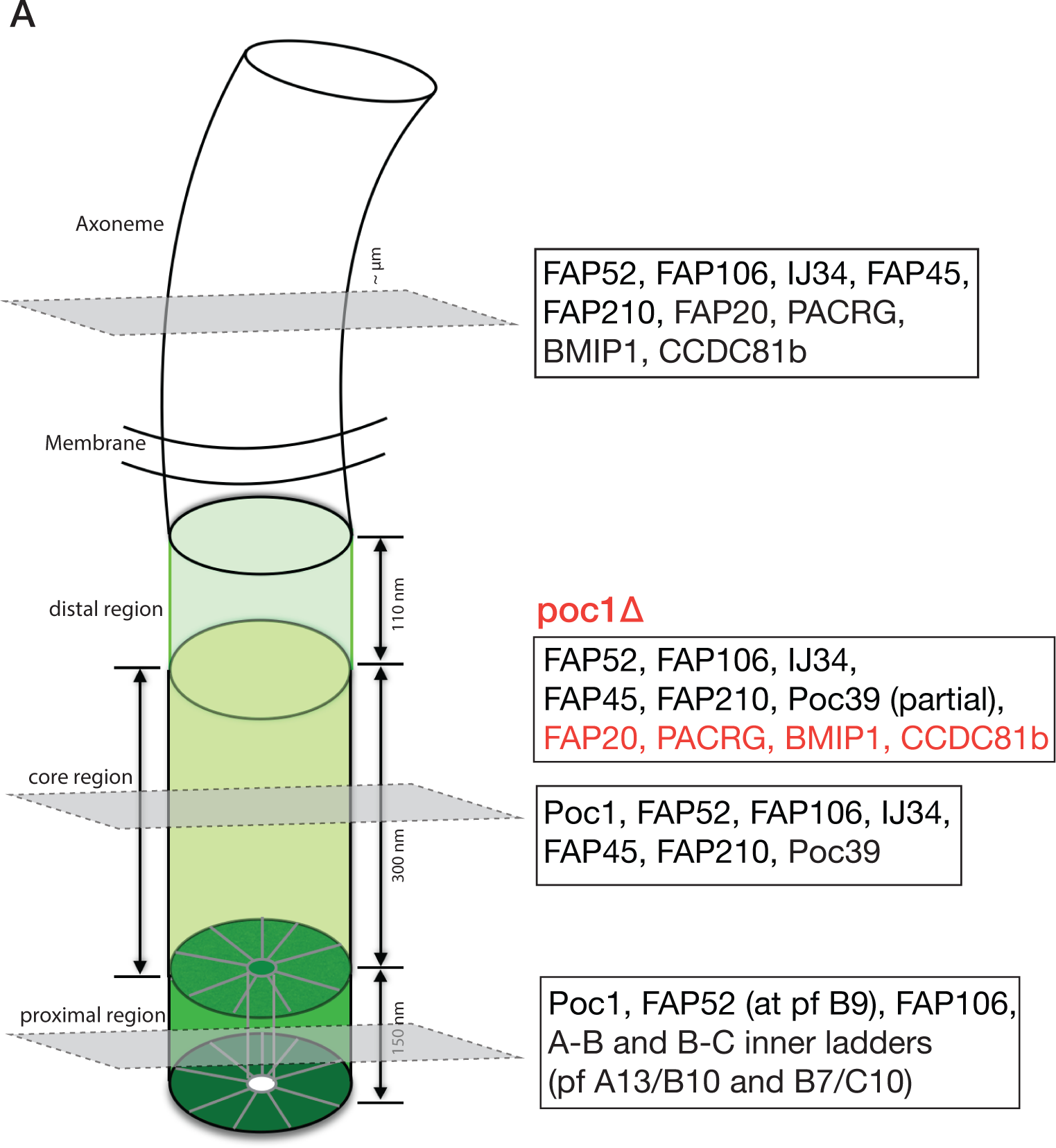
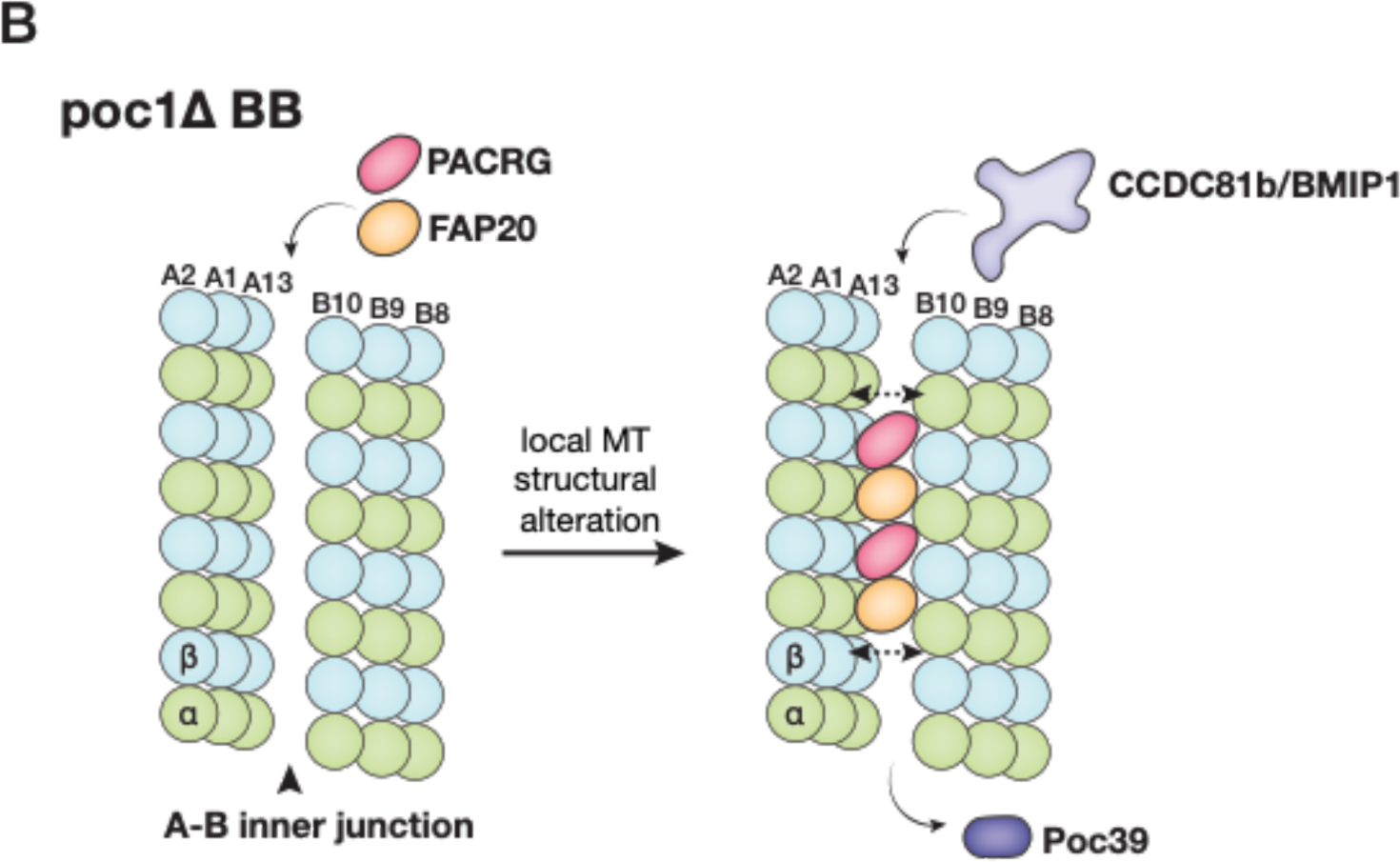
(A) Schematic illustration summarizing the inner junction components in three regions of the cilium. In poc1Δ, the axoneme-specific MIPs are in red; note that many BB components are partially bound in the mutant as the inner junction structure is disrupted without Poc1. (B) A model illustrates the poc1Δ BB A-B inner junction partially morphing into an axoneme-like architecture. The double-ended arrows indicate expansion of the inner junction gap as PACRG and FAP20 are incorporated in. This causes the disassociation of Poc39 and facilitates the binding of axonemal components, such as CCDC81b and BMIP1.

### MIPs shared by the BB and axoneme inner junction

Among the MIPs identified, FAP52 and FAP106 are present in all three A-B inner junctions. This suggests that both MIPs are recruited to the TMTs at the start of BB biogenesis and are continuously assembled throughout the cilium. Both FAP106 (ENKUR in vertebrates) and FAP52 (WDR16 or CFAP52 in vertebrates) are highly conserved proteins found in all high-resolution axonemal DMT structures to date, ranging from the protozoans including Trypanosome (Shimogawa et al., 2023), *Chlamydomonas* (Ma et al., 2019), and *Tetrahymena* (Kubo et al., 2023; Li et al., 2022), to higher eukaryotes (Zhou et al., 2023; Walton et al., 2022; Leung et al., 2023). Both proteins are associated with human ciliopathies (Ta-Shma et al., 2015; Sigg et al., 2017). Furthermore, FAP52 was implicated as a centrosome and BB component (Contreras and Hoyer-Fender, 2020; Hodges et al., 2010). This suggests essential roles for FAP52 and FAP106 in centriole/BB biogenesis and motile cilium assembly.

Three additional MIPs previously identified in the axoneme have also been found at the A-B inner junction in the BB core region. These are IJ34, FAP45 and FAP210. While IJ34 is unique to *Tetrahymena*, FAP45 and FAP210 are conserved across phyla and have been found in *Tetrahymena*, *Chlamydomonas*, bovine DMTs, and in mouse and human sperm axonemes (Ma et al., 2019; Gui et al., 2021; Kubo et al., 2023; Zhou et al., 2023; Leung et al., 2023; Chen et al., 2023). Both are filamentous MIPs binding end-to-end and exhibiting 48-nm periodicity. They bind to TMT and DMT similarly in the BB and axoneme. This confirms that the internal 48 nm periodicity is established in the BB core region and continues to the axoneme. Furthermore, a recent study on *T. brucei* axoneme revealed FAP106’s role as an “inner junction hub” critical for recruiting other MIPs, such as FAP45 and FAP210 (Shimogawa et al., 2023). Our identification of FAP106, FAP45 and FAP210 in the BB suggests that the hierarchical recruitment of MIPs found in the axoneme might also apply to the BB inner junction assembly.

The conservation of MIPs shared between the BB TMT and the axoneme DMT is likely not limited to the inner junction but extends to other parts of the A- and B-tubule (Ma et al., 2019), which we did not focus on in this work. Meanwhile, many previous phenotypical and functional analyses of MIPs have focused mainly on axonemes and flagella. However, their effects on the BBs have been grossly overlooked, mainly due to the relatively small size of the BB compared with the flagellum, structural polymorphisms, and the difficulty of biochemically isolating BBs. Given that many MIPs have structural roles in both BBs and flagellar axonemes, it is necessary to analyze both structures and their “connectome” to understand their impact on the cilium assembly and function. This is exemplified by a recent study on the axonemal protein CCDC39/40 (Brody et al., 2024), a heterodimer critical for recruiting other cilium components and dictating the axoneme’s 96-nm periodicity. Deletion of CCDC39/40 not only results in the loss of over 90 ciliary proteins and disjointed ciliary structural organization, but the impacts extend to several cilia-independent pathways, such as protein homeostasis and cell differentiation.

### The ciliary inner junction MIPs are in a well-defined longitudinal spatial location

The BB provides a template for forming a cilium. Initially, a BB emerges as a probasal body (pBB) or procentriole. To make a matured BB capable of templating the cilium, the pBB first elongates to about 150 nm, forming the proximal region. This is followed by the sequential assembly of the central core region, the distal region, the transition zone, and, eventually, the axoneme. Each region is signified by its distinct ultrastructural features, for example, the cartwheel in the pBB, the inner scaffold in the central core region, the termination of C-tubule at the distal region, the Dynein arms and the central pair MT in the motile cilium axoneme. These morphological differences imply composition changes throughout the cilium assembly under a well-defined sequential control. Our study on the inner junctions, presented in molecular details in three distinct regions along the longitudinal direction, agrees with this.

Interestingly, we found that FAP52 initially binds to pf B9 in the proximal region while an unidentified density, the A-B inner ladder, occupies pf B10 and bridges pf B10 and A13. FAP52 shifts location from pf B9 to pf B9/B10 at ∼130 nm. This coincides with the termination of the A-B inner ladder, suggesting that these two events might be coordinated. These are followed by the A-C linker’s termination and the inner scaffold’s emergence at ∼170 nm, marking the transition from the proximal to the core region. In the core region, additional MIPs are recruited to the A-B inner junction, including FAP45, FAP210, IJ34, and Poc39, which are unique to this region. In the axoneme region, Poc1 is replaced by PACRG and FAP20, and Poc39 is terminated while IJ34 remains. Coinciding with these composition changes, we observed MT structural alteration, likely to accommodate the changes, for example, variation in the local inter-protofilament lateral curvature and expansion of the lateral gap at the A-B inner junction in the axoneme.

The BB assembly initiates at its proximal end, where the TMT grows from the minus towards the plus direction. Our current study focused on the matured BBs that precluded assembly intermediates, such as pBBs. Thus, we cannot provide any temporal resolution for the assembly process nor know the relative timing when MIPs were incorporated into the BB. However, the spatial arrangement of the MIP observed in this study agrees well with the time-resolved gene expression profile in the ciliate *Stentor* (Sood et al., 2022). The study shows that the gene expressions involved in BB biogenesis and ciliary assembly are modular and can be described as a cascade wave of defined sets of genes. For example, POC1 is expressed before PACRG and FAP20, consistent with their sequential assembly in the BB and the axoneme. Similarly, a time-resolving RNA-seq study in *Tetrahymena* also indicates that expression of POC1 takes place before PACRG and FAP20 (Zhang et al., 2023). In summary, our observation of structures at three locations in the cilium is consistent with the notion that the assembly is a well-coordinated process under tight spatiotemporal control. However, a molecular understanding of the mechanism that governs this regulation is still incomplete.

### Axonemal proteins incorporated into BB inner junction in the absence of Poc1

Perhaps the most surprising finding in this study is identifying a set of axoneme-specific proteins, including PACRG and FAP20, in the poc1Δ mutant BB inner junction. The inner junction in the wild-type BB core region and the axoneme share several structural features. Both have FAP52, FAP106, IJ34, FAP45, and FAP210 that bind to the MT wall in the same periodicity (Fig 4B). However, the two inner junctions have marked differences. First, in the central core region, the A and B-tubules are crosslinked by Poc1, while in the axoneme, the place is filled by a PACRG/FAP20 filament. Second, there is a local geometry change. The lateral gap between pf A1 and pf B10 is wider in the axoneme compared to the BB core region (Fig 4F). Third, the MTs exhibit different local inter-pf curvature (Figure S4E, Table 2). Fourth, Poc39 is unique to the BB core region, while a set of MIPs, namely CCDC81b and BMIP1, are found only in the axoneme inner junction. The poc1Δ weakens the inner junction (Ruehle et al., 2024). The increased flexibility at the inner junction and the similarity of the local chemical environments to the axoneme will likely facilitate PACRG and FAP20 filling in the vacancy left by Poc1, leading to local structure changes. Once PACRG and FAP20 bind at the BB inner junction, it will promote recruiting additional PACRG and FAP20 and other axoneme components, such as CCDC81b and BMIP1. These structural changes result in a complete loss of Poc39 in the inner junction. Thus, in the poc1Δ BB, the A-B inner junction partially morphs from a BB architecture to that of an axoneme (Figure 6B).

Currently, we do not have a temporal resolution of these changes. Thus, we do not know the precise timing of PACRG/FAP20 incorporation in the poc1Δ BB inner junction. Nonetheless, given that the axoneme proteins PACRG/FAP20 are modularly expressed after the BB components (Sood et al., 2022; Zhang et al., 2023), it is likely that the incorporation takes place after the completion of BB assembly when PACRG/FAP20 congregate at the BB for cilium assembly (Yanagisawa et al., 2014). This implies that, in wild-type, the Poc1 assembled at the inner junction will prevent PACRG/FAP20 and other axonemal components from mistakenly being incorporated into the BB architecture, a regulatory mechanism used by the cell to maintain the spatial order of the components and ensure the structural integrity of the BB. In summary, the analysis of the inner junction structure in three regions, namely the proximal, the central core region of the BB and the axoneme, showcases the regulation of cilium biogenesis and provides structural insight into ciliary components, many of which can be linked to ciliopathies in humans.

## Online Methods

### Sample and EM grid preparations

The axoneme and the BB isolates from the *Tetrahymena* wild-type and poc1Δ strain have been described in detail (Li et al., 2022; Ruehle et al., 2024). The wild-type (B2086) *Tetrahymena* strain was obtained from the Tetrahymena Stock Center at Cornell University. The poc1Δ strain was generated as in (Pearson et al., 2009a).

The BB and axoneme EM grids were made at the University of Colorado Anschutz Medical Campus (Aurora, CO) or the University of California in Davis (Davis, CA), respectively. 200 mesh Quantifoil Cu or Cu/Rh 2/2 (Quantifoil, Inc) were used for all samples. A 4 µl sample mixed with 10 nm colloid gold beads coated with BSA was applied to the grid and plunge frozen in liquid ethane after various amounts of wait time (10-50 s.) using a Vitrobot (Thermo Fisher, Inc). The relative humidity in the Vitrobot chamber was 95%, the temperature was 22°C and the blot time was 0.5-1.0 s. The frozen grids were stored in liquid nitrogen and were transported to UCSF for data collection.

### Electron cryo-tomography data collection

Single-axis tilt series were collected for all samples on two field emission guns, 300 kV Titan Krios electron microscopes (Thermo Fisher, Inc) at UCSF. Each scope had a Bio-Quantum GIF energy filter and a post-GIF Gatan K2 or K3 Summit Direct Electron Detectors (Gatan, Inc.). The GIF slit width was set at 20 eV. SerialEM was used for tomography tilt series data collection (Mastronarde, 2005). The data were collected in the super-resolution and dose-fractionation mode. The nominal magnification was set at 33,000. The physical pixel size on recorded images was either 2.70 Å (for wild-type BB) or 2.65 Å (for poc1Δ mutant and wild-type axoneme). A dose rate was set at 20 electrons/pixel/second during exposure. A bi-directional scheme was used for collecting tilt series, starting from zero degrees, first tilted towards -60°, followed by a second half from +2° to +60°, in increments of 2° per tilt. The accumulated dose for each tilt series was limited to 80 electrons/Å^2^ on the sample.

### EM data processing and image analysis

For the tilt series alignment and tomogram reconstruction, the dose-fractionated movie at each tilt in the tilt series was corrected of motion and summed using MotionCor2 (Zheng et al., 2017). The tilt series were aligned using the gold beads as fiducials using IMOD and TomoAlign (Fernandez et al., 2018). The contrast transfer function for each tilt series was determined and corrected by TomoCTF (Fernández et al., 2006). The tomograms were reconstructed by TomoRec, which considered the beam-induced sample deformation during data collection (Fernandez et al., 2019). A total of 93 wild-type BB tomograms, 85 poc1Δ BB tomograms, and 51 wild-type axoneme tomograms were used for reconstruction.

For subtomogram alignment and averaging, the BBs or the axoneme were first identified in the 6xbinned tomograms. The center of TMT or DMT and their approximate orientation relative to the tilt axis were manually annotated in a Spider metadata file (Frank et al., 1996). The initial alignment and average were carried out in a 2x binned format (pixel 5.4 Å for the wild-type or 5.30 Å for poc1Δ mutant and axoneme). The longitudinal segment length of TMT or DMT in a subtomogram was limited to 24 nm and 50% overlapping with neighboring segments. In all cases, subtomogram alignments were carried out without using any external reference by a program MLTOMO implemented in the Xmipp software package (Scheres et al., 2009).

Since TMTs and DMTs are continuous filaments, a homemade program, RANSAC, was used to detect any alignment outliers and impose the continuity constraint on the neighboring segments after obtaining the initial alignment parameters. This corrected the misaligned subtomograms if only a few subtomograms within the same filament were misaligned. Otherwise, the entire filament was discarded without further processing. MLTOMO and Relion v3.1 or v4.0 were extensively used for the focused classification of the subtomograms (Bharat and Scheres, 2016; Zivanov et al., 2022). Customized soft-edge binary masks were used during classification to limit the analysis to the structure of interest. This was critical for determining the correct periodicity of the MIPs and identifying structural defects or heterogeneity in the structure. Once the correct periodicity was found, these out-of-register subtomograms were re-centered and re-extracted. This was followed by combining all subtomograms for the next round of refinement. Previously, we found the MIPs at the inner junction in the core region of BB having 16 nm periodicity. It was based on analysis limited to a small cross-section region, including pfs B9, B10, A1, and A13. We expanded the study to a larger area, including more pfs. This led us to the identification of fMIPs FAP45 and FAP210 in the BB central core region. Both have a 48-nm periodicity, the same as in the axoneme. The final refinements, focusing on the inner junctions, were implemented using a workflow in Relion v4.0 (Zivanov et al., 2022). The overall resolutions (Figure S1, Table 1) are based on the Fourier Shell Correlation (FSC) cutoff at 0.143 (Rosenthal and Henderson, 2003; Scheres and Chen, 2012).

### Model building and analysis

The pseudo-atomic models for the inner junctions in different regions of the BB and the axoneme were built based on the subtomograms averaged density maps described in this work. UCSF ChimeraX (Pettersen et al., 2021) was used for model building. The atomic models for individual MIP were based on previously published data (PDB ID 8G2Z) (Kubo et al., 2023) or models predicted by AlphaFold2 (Jumper et al., 2021), which was installed and run locally on the UCSF Wynton HPC, or AlphaFold3 (Abramson et al., 2024) via the AlphaFold server. The models for individual MIP or α/β tubulins were fit into the subtomogram averaging density maps, all in subnanometer resolution, using the “FitMap” command in UCSF ChimeraX. Two constraints were applied when fitting the MIPs whose atomic models were available from the previous axoneme structure, the location and the tertiary fold of a specific MIP. For the AlphaFold3 predicted structure, the ChimeraX built-in function “AlphaFold error plot” was used to color the predicted model based on its Predicted Aligned Error (PAE) score provided by the AlphaFold server. UCSF ChimeraX was also used for visualization and for recording images and videos.

To analyze the microtubule inter-protofilament angle and curvature, the UCSF ChimeraX command “matchmaker” was first used to bring two reference tubulin models to the two targets in the neighboring protofilaments. This was followed by the “measure rotation” command to find the rotation angle between the two reference models. The output is the inter-protofilament angle, a local MT lateral curvature measurement.

To measure the difference of inner junction width between pfs A1 and B10 at the regions of BB or axoneme, once the MT models were built, ChimeraX command “matchmaker” was used to bring the two models to the same reference; in this case, the pf A1 served as the reference. This is followed by a second command, “rmsd #Model1@ca to #Model2@ca”. This provided the root-mean-square deviation (RMSD) between the two molecules in pf B10, measuring the width change between the inner junctions.

### GFP-Poc39 Tagging and Immunofluorescence microscopy

GFP-Poc39 is expressed from the Mtt1 Cadmium inducible promoter using the pBS.MTT-GFP vector (gift of Douglas Chalker, Washington University). The POC39 gene was cloned in frame following the GFP reading frame and cells were transformed with linearized vector using the BioRad PDS-1000/He Biolistic Particle Delivery System and selected with 8.0 ug/ml cycloheximide in modified Neff growth media (Bruns and Cassidy-Hanley, 1999). Three independent cell lines were obtained that all showed similar localization. Cells were grown overnight in 0.3 ug/ml CdCl2 to induce gfp-Poc39 expression. Cells were collected by centrifugation and resuspended in 10 mM TRIS for 2h prior to fixation in 3.2% formaldehyde and 2.4% TritonX-100 in PHEM buffer (60 mM PIPES, 25 mM HEPES, 10 mM EGTA, 2 mM MgCl2). Immunofluorescence to show co-localization with centrin in basal bodies was performed as described previously using a rabbit polyclonal anti-Cen1 antibody (1:1000) (Stemm-Wolf et al., 2005) and AlexaFluor594 conjugated goat anti-rabbit secondary antibody (1:1000) (Life Technologies). Images were obtained using a Nikon Eclipse Ti widefield fluorescence microscope. Z-stacks were collected at 0.5 μm intervals and images were deconvolved using the low noise setting on the Nikon NIS-Elements software. Stacks were projected in ImageJ to create images shown.

### Tetrahymena Gene Expression Profile

The gene expression profiles were obtained via the Tetrahymena Genome Database website (https://tet.ciliate.org/index.php). The following *Tetrahymena* genes are used for the query: POC1 (TTHERM_01308010), FAP20 (TTHERM_00418580), PACRGA (TTHERM_00446290), PACRGB (TTHERM_00499570), PACRGC (TTHERM_00499310).

## Abbreviations

BB: basal body
TMT: triplet microtubule
DMT: doublet microtubule

## Acknowledgments

We thank Brian Wimberly, Dave Farrell, Peter Van Blerkom, Eduardo Romero Camacho (University of Colorado School of Medicine, Denver), and Fei Guo (University of California, Davis) for help with screening and preparing EM grids; Annie Rhee (University of California, Davis) for technical assistance. We thank David Bulkley and Eric Tse (UCSF) for assistance on tomography data collection, the Wynton HPC team (UCSF) for supporting the computational infrastructure, Tom Goddard (UCSF) for help with AlphaFold2 and UCSF ChimeraX software. The UC Davis BioEM Facility is supported by user fees, the Department of Molecular and Cellular Biology, the College of Biosciences, the Office of Research and the Provost’s Office. This work is supported in part by NIH grants GM127571 (M.E.W.), GM118099 (D.A.A.), GM140813 (C.G.P), and by the Spanish AEI/FEDER (PID2022-139071NB-I00) (J.-J.F.).

The EM structures have been deposited in the EMDB with the following accession numbers: EMD-46437, EMD-46438, EMD-46439, EMD-46440, EMD-46441, EMD-46442, EMD-46443.

---

**Figure S1.**
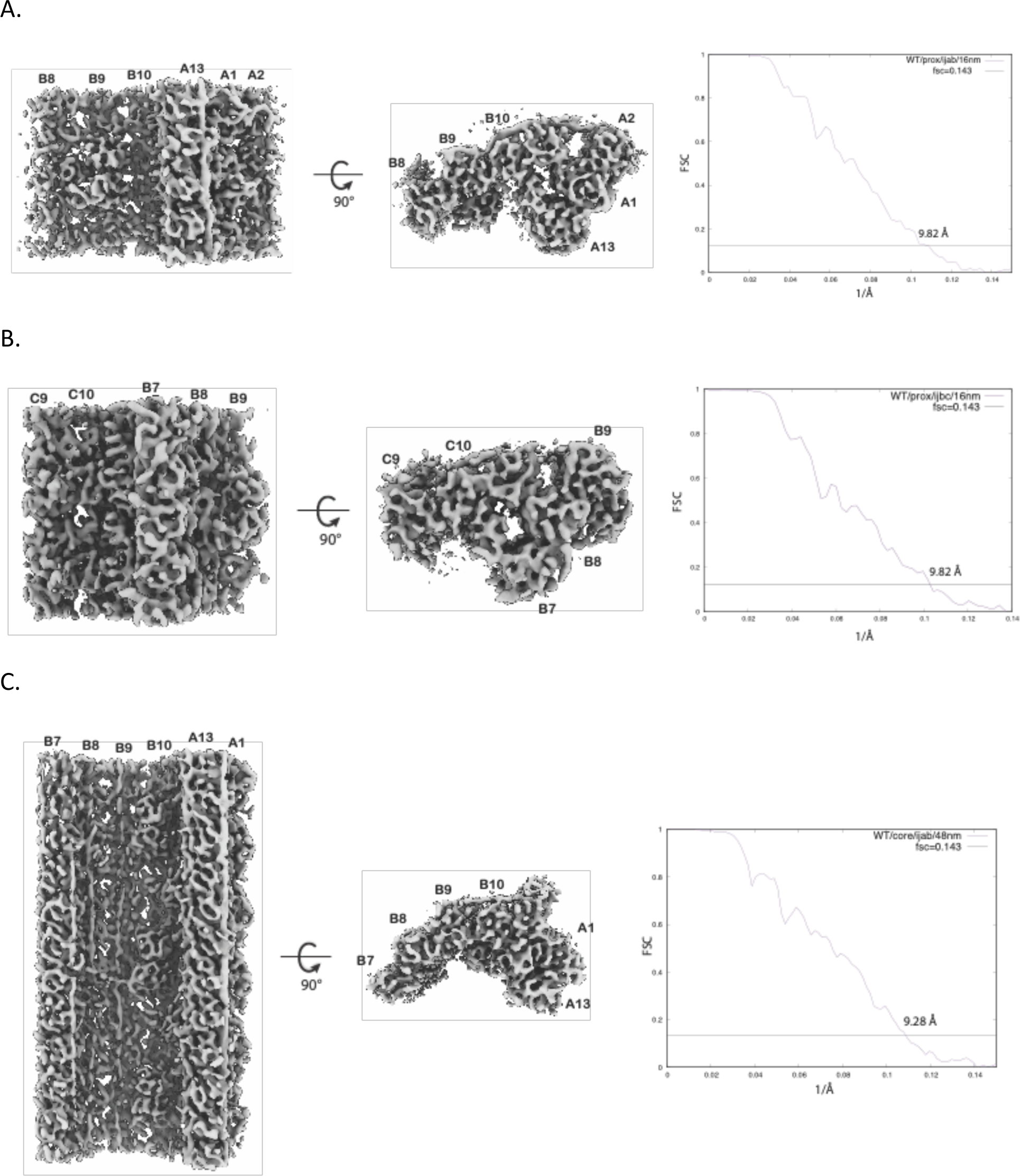
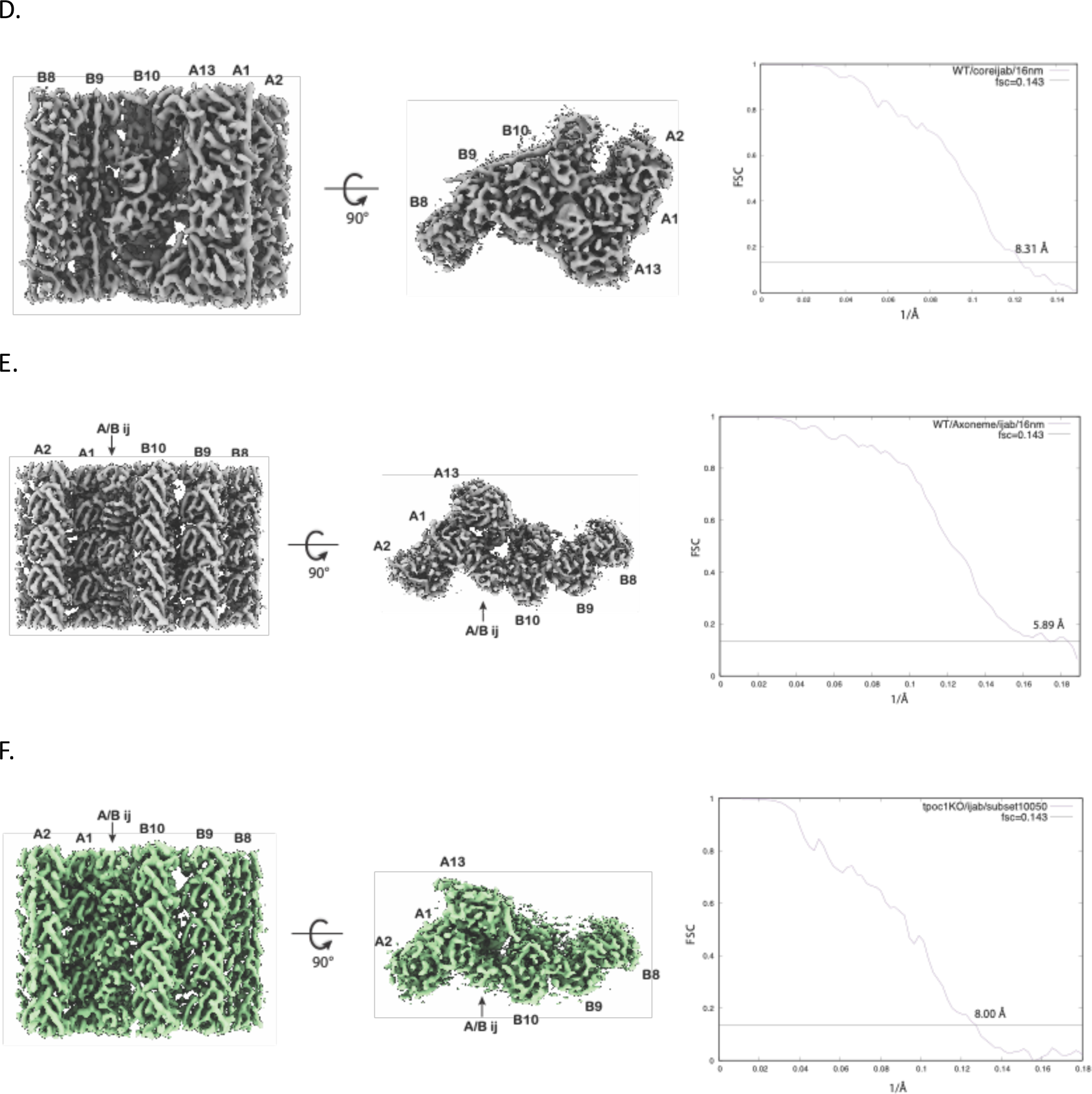
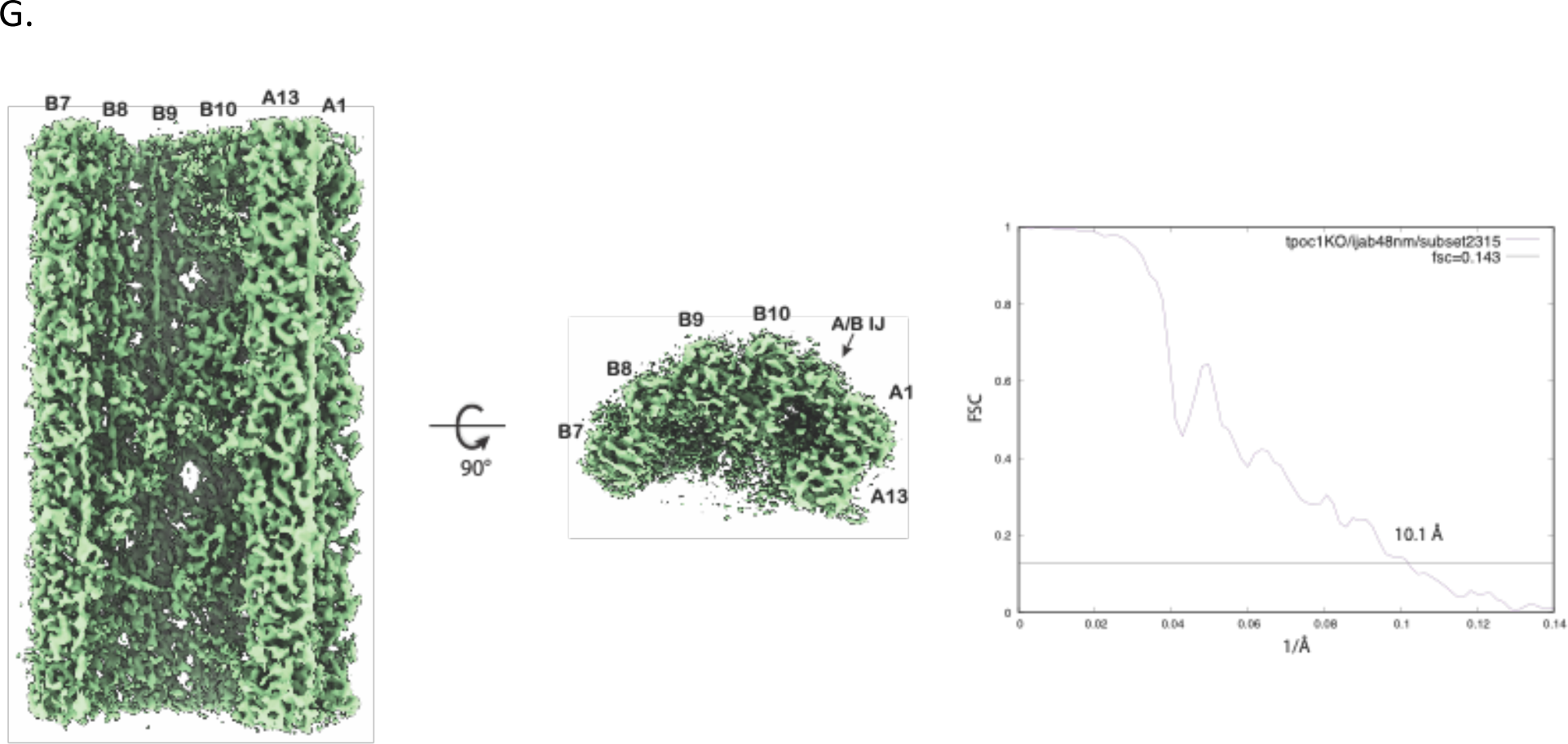
*Related to* Figures 1*, 2, 4, 5, S5*. Assessing resolution of subtomogram averages by Fourier shell correlation (FSC). The structures and their Fourier shell correlation as a function of resolution (1/Å) as reported in Table 1. (A) A 16-nm repeat of the A-B inner junction from the proximal region of BB (wild-type). (B) A 16-nm repeat of the B-C inner junction from the proximal region of BB (wild-type). (C) A 48-nm repeat of the A-B inner junction from the central core region of BB (wild-type). (D) A 16-nm repeat of the A-B inner junction from the central core region of BB (wild-type). (E) A 16-nm repeat of the A-B inner junction from the axoneme (wild-type). (F) A 16-nm repeat of the A-B inner junction from a subset (Class 3) of the central core region of poc1Δ BB. (G) A 48-nm repeat of the A-B inner junction from a subset (Class 3) of the central core region of poc1Δ BB.

**Figure S2.**
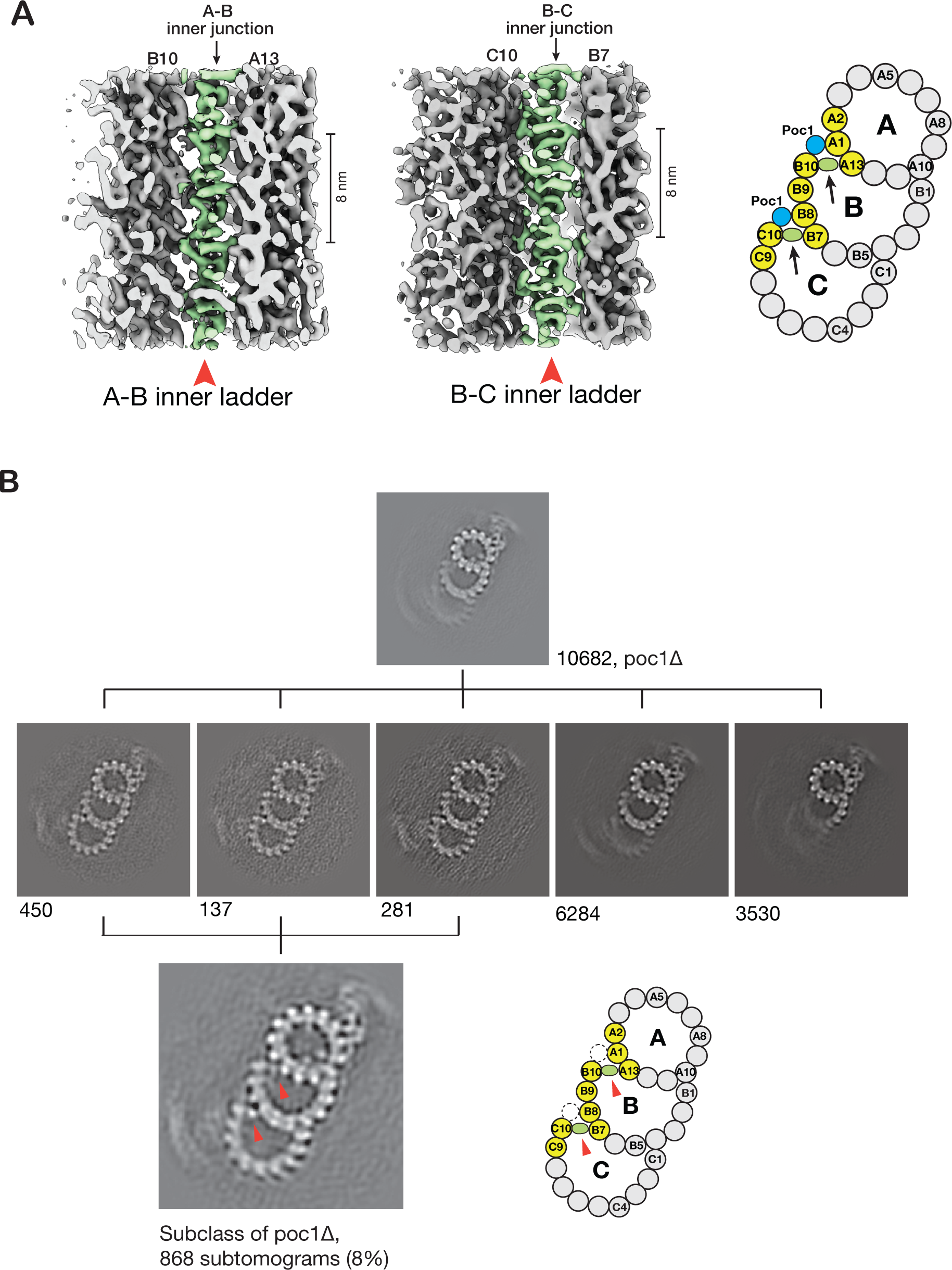
*Related to* Figure 2. Two unidentified proteins at the inner junctions in the proximal region. (A) The density maps of the A-B and B-C inner junctions in the proximal region of BB. The two unidentified proteins, namely the A-B inner ladder and B-C inner ladder crosslinking pfs A13-B10 or B7-C10, respectively, are highlighted in green and indicated by red arrowheads. On the right, a schematic diagram shows the location of the above two maps in the TMT. Black arrows indicate the viewing directions. (B) 3D Classification of the subtomograms from the proximal region of poc1Δ TMT. A small fraction of the dataset (8%) shows complete TMT, where the two unidentified proteins, the A-B inner ladder and B-C inner ladder indicated by red arrowheads, remain in the inner junctions. The number of subtomograms in each class is shown. The dashed circles in the cartoon indicate the location of Poc1 in the wild-type.

**Figure S3.**
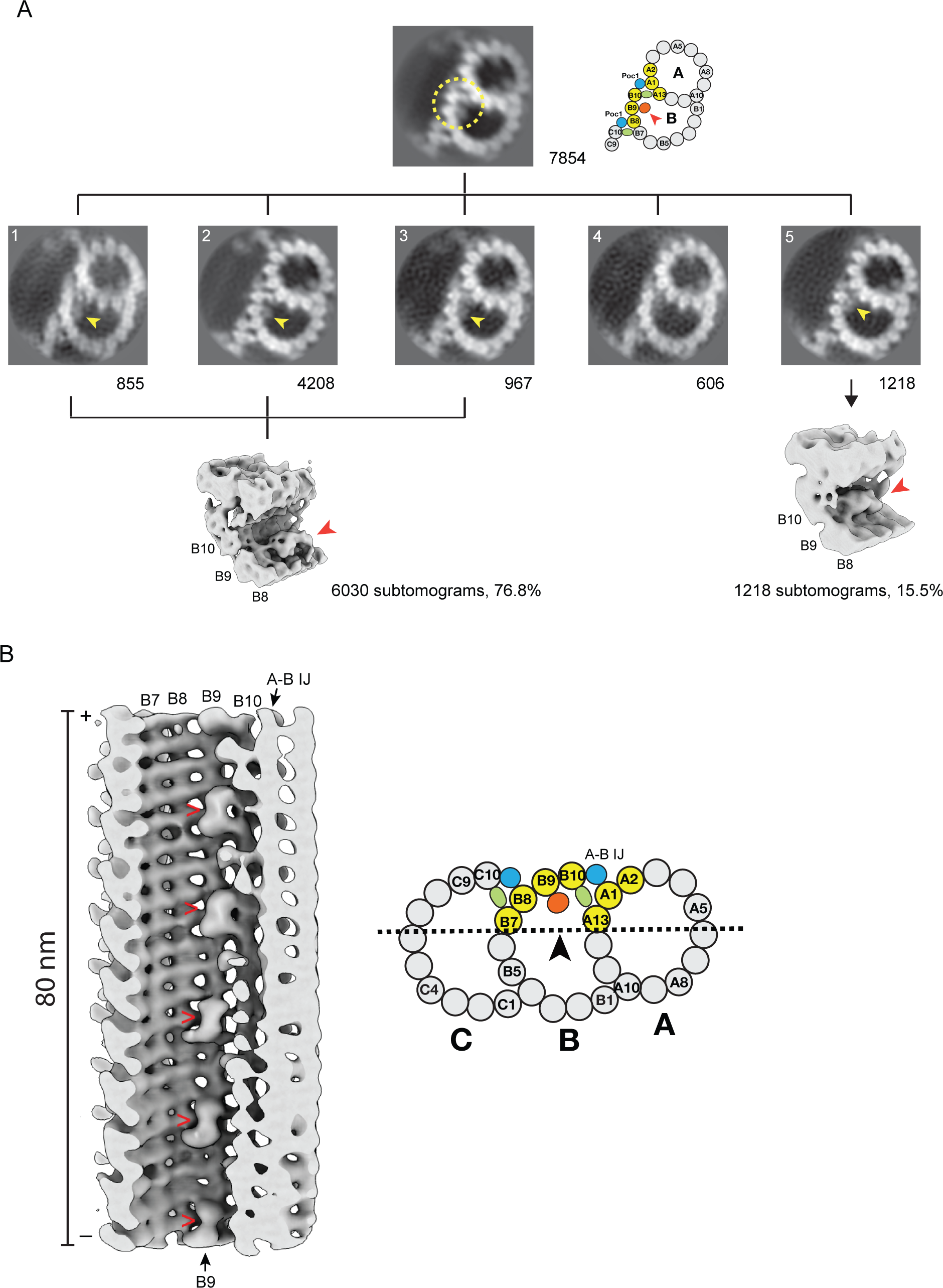
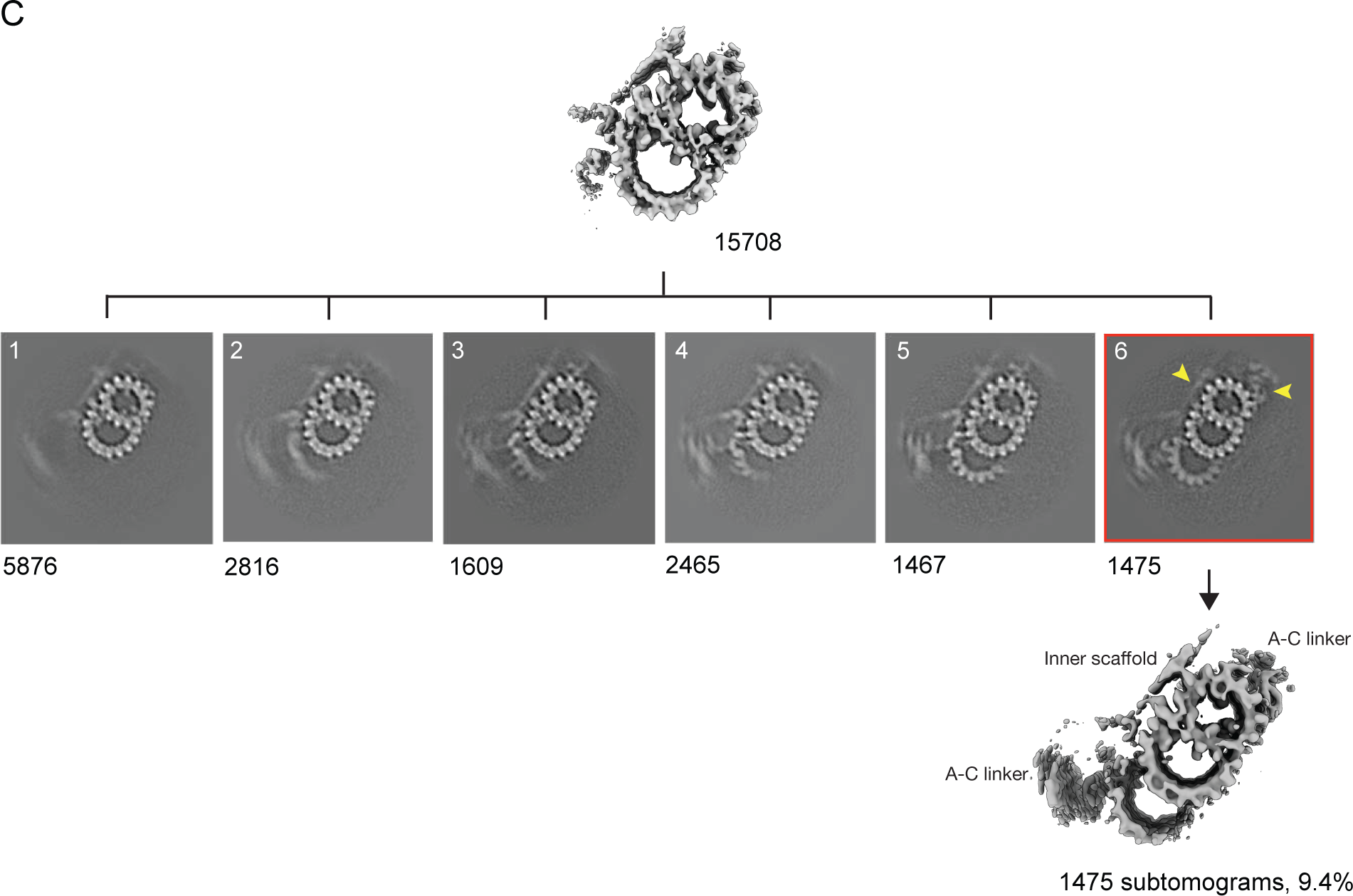
*Related to* Figure 3. (A) Focused 3D classification on the subtomograms from the proximal region of the BB. A yellow dashed circle indicates the focused area centered on the inner junction. Yellow or red arrowheads indicate the locations of FAP52 in the class averages. (B) A Longitudinal cross-section of the A-B inner junction showing the FAP52, indicated with red arrowheads, shifts binding from pf B9 to pf 9/10. A schematic illustration of the TMT is on the right. A dashed line and arrow indicate the cross-section and viewing direction of the structure on the left. An orange circle represents FAP52. (C) Focused 3D classification on the subtomograms from the central core region of the BB. The Class 6 is highlighted with a red frame. In this class, the A-C linker and the inner scaffold are indicated by yellow arrowheads. The number of subtomograms in each group is provided.

**Figure S4.**
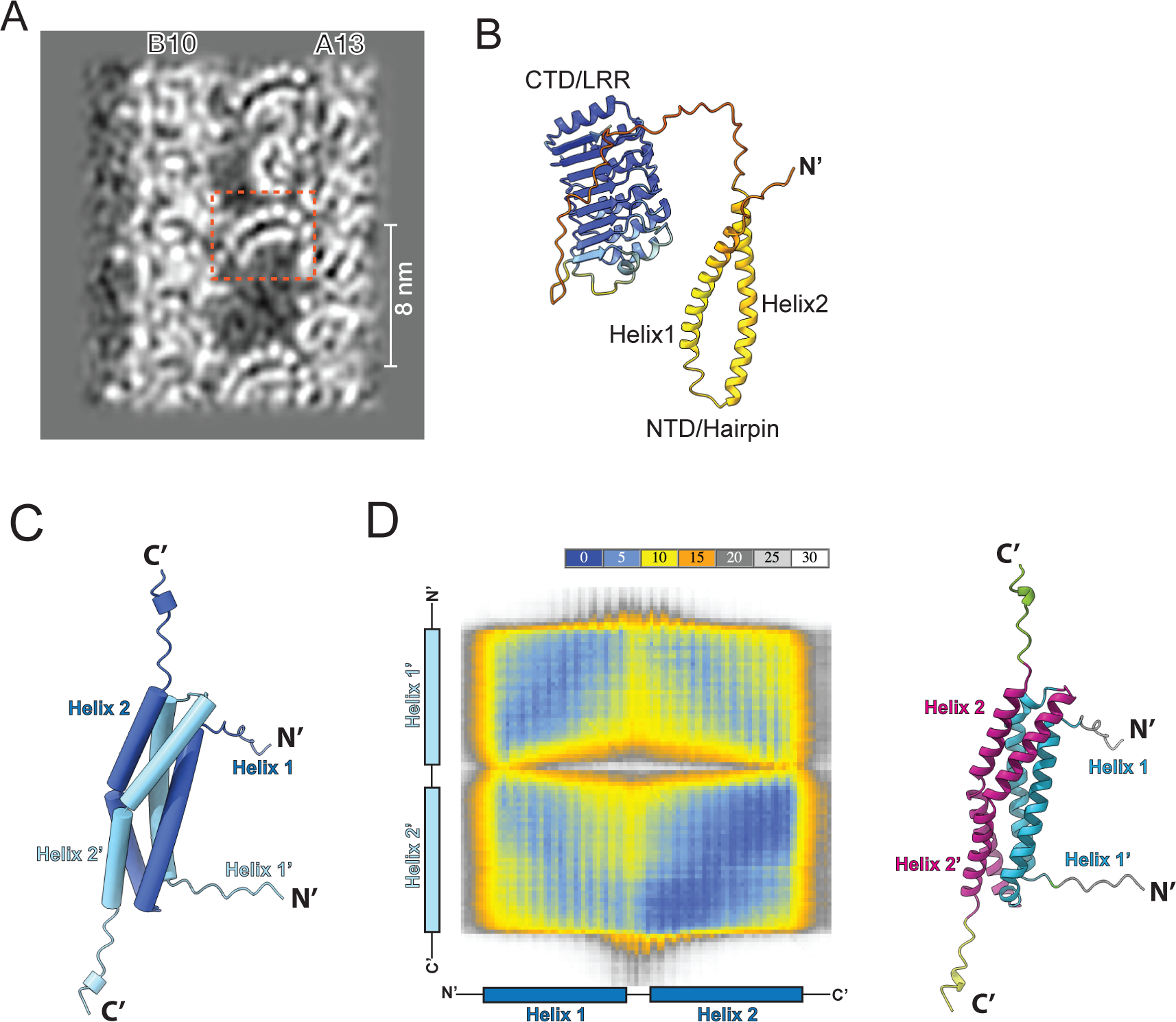
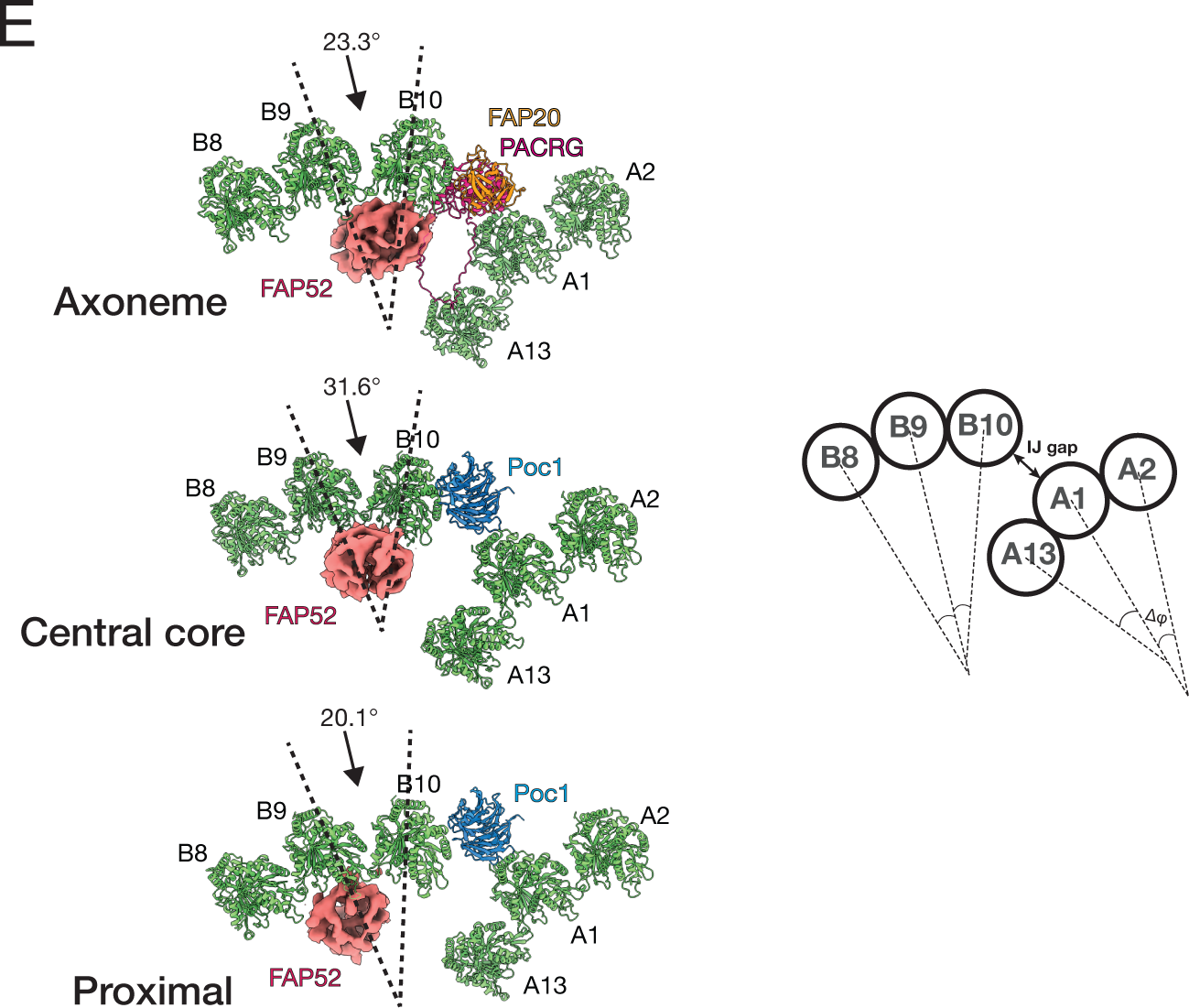
*Related to* Figure 4. (A) A cross-section slice of the density map shows an LRR motif highlighted in a red dashed-line square. (B) An AlphaFold2 predicted protein structure (UniProt Q22N53) was identified previously in the BB proteome. The protein is composed of a N-terminal α-helix hairpin (NTD) and a C-terminal LRR motif (CTD). The structure is colored based on the prediction confidence score (pLDDT: the predicted local distance difference test). The high confidence is in dark blue, while the low confidence is in yellow or orange. (C) An AlphaFold3 predicted 4-helix bundle formed by dimerizing two NTD/hairpins. Two copies of the NTD hairpin form an anti-parallel dimer. One monomer is in dark blue (Helix 1 and Helix 2) and the other monomer is in light blue (Helix 1’ and Helix 2’). The dimer forms a right-handed 4-helix bundle. (D) The predicted aligned error (PAE) plot provides inter-domain packing confidence scores. The dark and light blue imply the prediction with high confidence, while the grey and white indicate low confidence in the interaction. On the right, a ChimeraX-adapted color scheme is used where the 4-helix bundle is colored based on the PAE potential interaction score. The two N-terminal helices (Helix 1, Helix 1’) are in cyan, and the two C-terminal helices (Helix 2 and Helix 2’) are in magenta, indicating that the interaction between Helix 1 and Helix 1’ (cyan), Helix 2 and Helix 2’ (magenta) are with high confidence. (E) Inter-protofilament angle measurement. Left: the angles between pfs B9 and B10 are measured at the three regions, showing varying local curvature. The FAP52 and Poc1 or FAP20/PACRG are shown as reference points. Right: a schematic diagram depicts the inter-protofilament angles at the A-B inner junction. More complete measurements are in Table 2.

**Figure S5.**
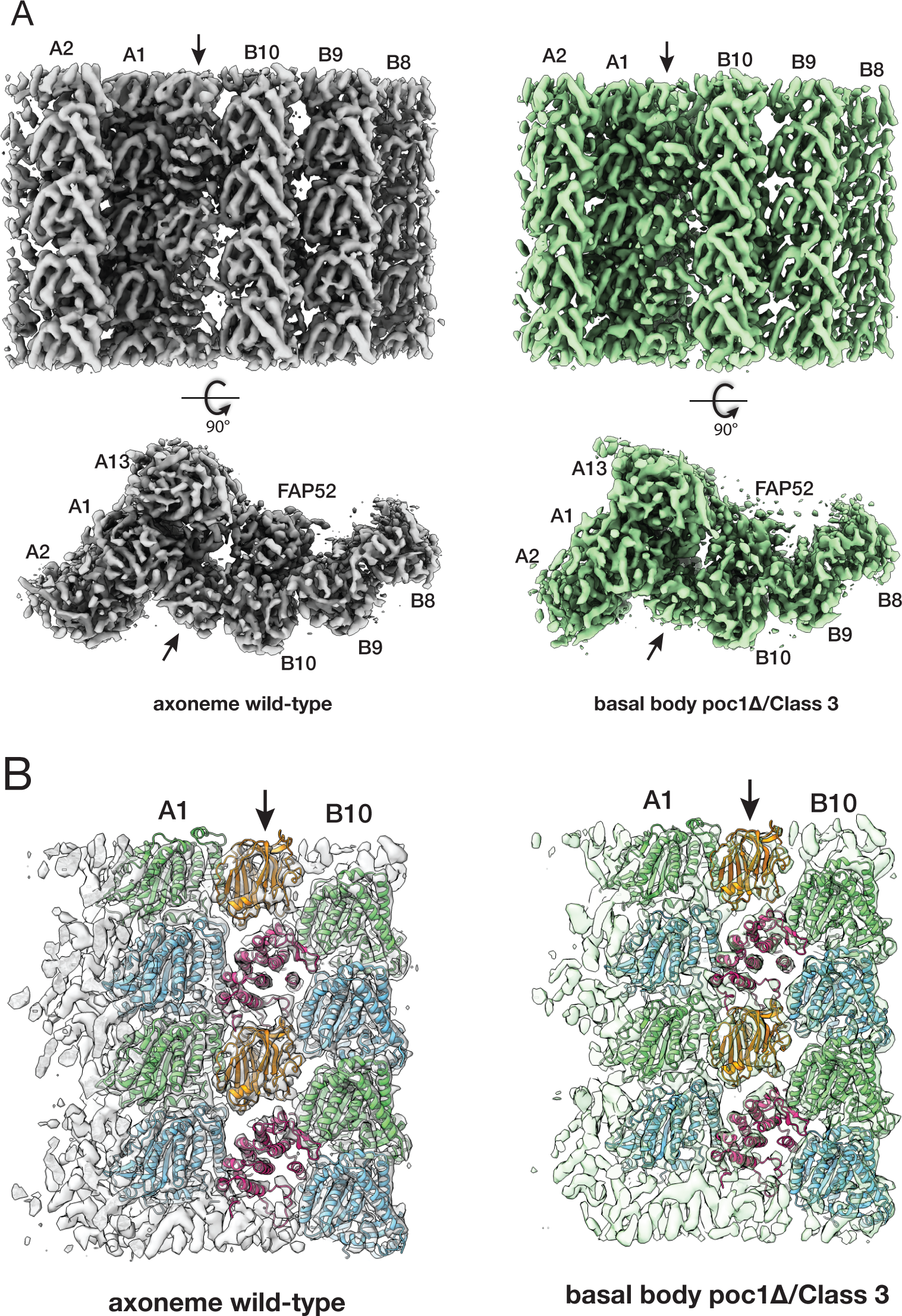
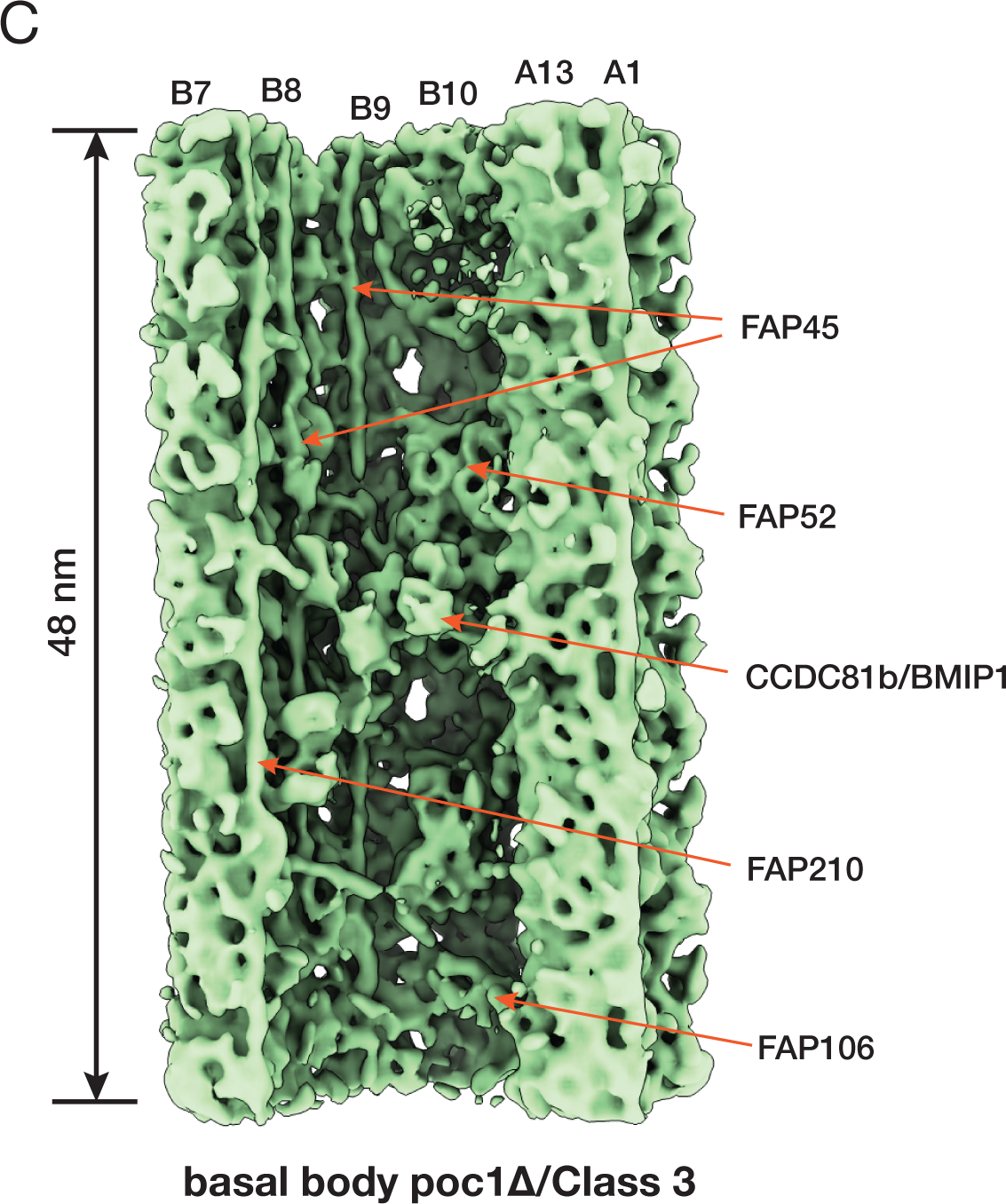
*Related to* Figure 5. Comparing the inner junctions between the wild-type axoneme and the Class 3 from the poc1Δ BBs. (A) Comparing the two structures in two orthogonal views, the wild-type axoneme is in grey, and the poc1Δ BB is in green. (B) Fitting the molecular models into the density maps in (A). The α/β tubulins are in light green and blue, FAP20 is in orange, and PACRG is in purple. The arrows indicate the A-B inner junctions. (C) A 48-nm repeat average from a subset of poc1Δ BB (Class 3) shows nearly identical structure to the wild-type axoneme inner junction. This is the same subset as shown in Figure 5E, but here the longitudinal length is extended to 48 nm. For clarity, the map is low-pass filtered to 12 Å.

